# Natural genetic transformation of the wheat rhizosphere microbial community through DNA inoculations

**DOI:** 10.64898/2025.12.18.695157

**Authors:** Charlotte Giard-Laliberté, Julien Tremblay, Hamed Azarbad, Luke Bainard, Étienne Yergeau

**Author notes:** Correspondance: Etienne Yergeau, Institut national de la recherche scientifique, Centre Armand-Frappier Santé Biotechnologie, 531 boul. des Prairies, Laval (Québec), Canada, H7V 1B7.

## Abstract

Rhizosphere microorganisms are known to be able to modify the plant’s ability to resist abiotic stresses. It is, however, difficult to modify microbial communities to improve plant phenotypes. Here we tested if a rhizosphere microbial community from a water-stress naïve soil could be modified by adding DNA extracted from soils with a water stress history. Six-week-old wheat plants growing under low or high-water availabilities were inoculated with DNA extracted from soils with contrasting long-term histories of water availability – one continuously and the other intermittently exposed to water deficit. The fate of the inoculated DNA in the rhizosphere microbial communities was assessed by shotgun metagenomics. Putatively transferred inoculum genes were disproportionately found in the *Acidobacteria* and *Bacteroidetes* and belonged to functional category such as antibiotics, biofilm, and carboxylates metabolism, among others. These functional categories were shared by pre-inoculation laterally transferred genes in the recipient soil, highlighting their usefulness for life in soil. The “continuous” inoculum reduced the stress levels of wheat under reduced soil water content, suggesting that the natural genetic transformation of the rhizosphere community can feedback to the plant. Altogether, we are providing evidence for an ecological mechanism that could be harnessed to modify plant-associated microbial communities and help plants sustain water stress.

## Introduction

Although there are numerous reports on the genetic, molecular and physiological basis of how plants respond to drought stress [1], we do not know the exact nature of plant adaptation to high stress. When we subjected wheat to water stress, most of the transcriptomic response came from the rhizosphere bacteria [2], suggesting that management or engineering of the plant associated microbiota [3, 4] could help plant adapt to water stress. Many specific bacteria of the phylum *Actinobacteria* and *Proteobacteria* can improve plant tolerance to drought- or salinity-related stresses [5–8]. Fungal endophytes can also improve plant performance under abiotic stress [9–11]. Mycorrhizal fungi can improve water use efficiency and reduce drought stress in wheat [12], oat [13], and corn [14]. Interestingly, endophytic and rhizospheric microorganisms isolated from environments prone to drought confer plants with a better resistance to drought [11, 15, 16].

Many mechanisms are involved in the enhancement of plant drought tolerance by microbes. These include modulation of plant drought stress genes [17], reduction of the stress hormone ethylene levels through degradation of its precursor 1-aminocyclopropane-1-carboxylic acid (ACC) by the bacterial enzyme ACC deaminase [5, 6], stimulation of the expression of plant genes related to osmolytes and osmoprotectants by bacterial volatile organic compounds [18] and modulation of the plant epigenetics response to drought [19]. To harness these microbial mechanisms and improve plant drought tolerance, we need to shape the plant microbial communities.

There are four major mechanisms by which a community can change: 1) migration, 2) selection, 3) speciation, and 4) drift [20]. Most efforts to shape plant and soil microbial communities to date have focussed on the “migration” mechanism, through inoculations of selected microorganisms. This is, however, not the most ecologically sound approach in the context of these hyper diversified communities [4]. Indeed, displacing an already established highly diversified community is difficult, due to priority effects and competition [21, 22]. Even when we inoculated complex microbial communities extracted from water stressed soils, the bulk of the wheat rhizosphere microbial response to water stress was through “selection” – i.e. reduction or amplification of their relative abundance – of the microorganisms already in the rhizosphere [23]. Only 2 to 5% of the rhizospheric bacteria of stressed plants had “migrated” from the bulk soil and inoculum [23], suggesting that this is not the main mechanism involved in microbial community response to water stress. In another study, we inoculated 25 highly osmotolerant species – containing many *Firmicutes*, *Bacteroidetes* and *Actinobacteria* –, which increased wheat fresh weight under low soil water content [16]. However, the inoculation also modified the abundance of other rhizospheric bacteria, many of which could have helped wheat withstand water stress [16]. Clearly, “migration” is not a panacea, and we should study other community modification mechanisms.

Here, we aimed at testing lateral gene transfer (LGT) as a “speciation” mechanism to rapidly modify the wheat rhizospheric microbial community under water stress. Since drought resistance is clearly a polygenic trait (i.e. based on multiple genes, such as genes related to osmolytes, antioxidants, biofilm formation, and plant hormones) [24], we used an approach based on natural transformation with DNA from highly diversified drought-adapted soil microbial communities rather than focusing on specific genes. Many bacteria and archaea are naturally competent and not only take up DNA pieces from the environment but integrate them in their genome, leading to natural transformation [25]. Although the natural transformation of *Bacillus* and *Pseudomonas* in soil/sediments is well known [26–28], some studies reported that naturally transformable rhizospheric bacteria are rare [29, 30]. However, previous studies were performed *in vitro* or in the absence of stressors, which are known to induce competence [25]. We first hypothesized that wheat rhizosphere microbial communities under low water availability would integrate more genes than communities under high water availability. We also hypothesized that, if it occurred, the transfer of genetic material would be biased toward genes that confer traits important to survival and reproduction in the recipient ecological niche.

## Material and methods

### Experimental design and sampling

To test our hypotheses, we set-up a two-level factorial greenhouse pot experiment with two soil water content (SWC, 15 or 50% of the soil water holding capacity, SWHC) x three inoculation treatments (inoculation with DNA extracted from continuous or intermittent Saskatchewan wheat field soils, or water as a non-inoculated control). The factorial combinations were replicated six times in a randomized complete block design, resulting in 36 pots. We filled each pot (1,000 cm^3^) with 1.5 kg (dry weight) of soil from our experimental field into which we sowed 8 seeds of *Triticum aestivum* cv. AC Nass, a *Fusarium*-resistant wheat cultivar developed for Eastern Canada (not bred for drought resistance). For the first four weeks of the experiment, we maintained all the pots at 50% SWHC, after which we decreased the water content of half the pots to 15% SWHC (LW: low water content treatment) while we maintained the other half at 50% SWHC (HW: high water content treatment). Two weeks after adjusting the water content, we inoculated the pots with 50 mL of one of the two inoculums (continuous or intermittent) or 50 mL of distilled water.

### Potting soil and DNA inoculum preparation

We collected potting soil from our experimental field at the Institut national de la recherche scientifique (Laval, QC, Canada) in May 2017. The field was ploughed for the first time in 2016 with no agricultural crops cultivated in the area for over 20 years. The region of Montreal, where the field is located, is not usually subjected to significant drought event, so we considered these soils as water-stress naïve. Soil analyses were carried out in May 2016 by Maxxam (Montréal, QC) and revealed a total P concentration of 1,300 mg/kg, a total K concentration of 1,110 mg/kg, a pH of 7.30 and a total N concentration (Kjeldahl extraction) of 3,000 mg/kg. All the soil was dried, sieved with a 2 mm mesh sieve and homogenized. The soils used for the DNA inocula came from two adjacent Agriculture and Agri-food Canada experimental fields in Swift Current, SK, Canada. These fields have been under a continuous wheat-fallow rotation to test newly developed wheat varieties for their resistance to water stress. Since 1981, the two fields have been managed similarly, except that one was irrigated every second year (during the wheat phase of the rotation) [23, 31, 32]. The climate in this region is considered semiarid with average annual precipitation of 345 mm, meaning that the wheat field under ambient conditions was continuously exposed to water stress for almost 40 years, whereas the irrigated wheat field was intermittently exposed to water stress. We sampled both soils in April 2017 and we sieved them with a 2 mm mesh size sieve. For each of the two soils, we did 80 large-scale DNA extractions from 15 g of soil using a scaled-up homemade phenol-chloroform protocol [31]. We pooled and diluted the extractions before the inoculations, from which we inoculated 50 ml at a concentration of 17.6 ng/µl (intermittent inoculum) or 17.7 ng/µl (continuous inoculum) directly in the rhizosphere, for a total of 880-885 µg of DNA inoculated. This resulted in each of the 24 inoculated pots receiving DNA extracted from an equivalent of 100 g of soil (for each soil type, 80 extractions from 15 g of soil divided in 12 pots = 100 g per pot). Before the inoculation, we subsampled each of the two inoculums for metagenome sequencing.

### Sampling

Three weeks after inoculation, we terminated and sampled the experiment. We took half a gram of the flag leaf, flash-froze it immediately with liquid nitrogen and stored it at -80°C before enzymatic measurements. To measure fresh and dry biomass and plant and leaf water content, we sampled, weighed, and dried at 80°C the remaining aboveground biomass. We uprooted the plants, and the soil still attached to the roots after vigorous shaking was considered rhizosphere soil, which we stored at -20°C for microbial analysis.

### Enzymatic assays

We measured superoxide dismutase (SOD) and catalase (CAT) activities as an indicator of plant water stress [33]. For SOD, we first homogenized the flag leaf in liquid nitrogen using a mortar and pestle, then mixed it with 1 mL of the extraction buffer composed of 100 mM phosphate buffer (pH 7.5), 1 mM EDTA, 5% W/V PVPP and 1 mM of ascorbic acid [34], as well as 1 µL of protease cocktail inhibitor (Sigma P9599). We vortexed the resulting mix and centrifuged it at 10,000 × g for 20 minutes at 4°C. We stored the supernatant at -80°C before quantifying SOD activity with an assay kit (Cayman, Michigan, USA). We normalized the results according to the total protein content measured with the Bradford assay method [35] and the Bio-Rad protein assay dye kit (Bio-Rad Laboratories, Hercules, CA, USA). We measured the CAT activity as previously described [23].

### DNA extraction and metagenomic sequencing

We extracted DNA from 500 mg of rhizosphere soil using the Qiagen DNeasy PowerLyzer PowerSoil Kit. Extractions yielded approximately 50 ng/µl on average, for a total of 5 µg of DNA. We also extracted DNA from the Saskatchewan soil used to produce the DNA inocula. The 40 DNA extracts (36 from the greenhouse experiment, 2 from the Saskatchewan soils, and 2 subsamples from the two DNA inocula) were then sent to the Centre d’expertise et de services Genome Quebec (Montreal, Canada) for Illumina TruSeq library preparation and Illumina HiSeq 4000 PE100 sequencing.

### Bioinformatic analyses

The metagenome sequencing of the 40 samples resulted in 291 Gbp which we processed through our metagenomics bioinformatics pipeline [36], as previously described [2]. We did two different assemblies: 1) using the two inoculum samples and 2) using the rhizosphere samples without the inoculum samples. We then mapped all the reads on the assemblies, resulting in an “inoculum” and a “no-inoculum” dataset. We used the “non-inoculum” dataset for general community and functional analyses of the rhizosphere samples and for the WAAFLE procedure. We used the “inoculum” dataset to find putatively transferred genes from the inoculum, as it excluded the rhizosphere reads that did not map to the inoculum contigs.

### Statistical analyses

We performed all analyses in R (v. 4.5.1), except for the WAAFLE program [37] for which we used the default parameters (https://github.com/biobakery/waafle). The R scripts used to manipulate data, perform statistical analyses and generate tables and figures are available on GitHub (https://github.com/le-labo-yergeau/DNA_Inoculation). To identify putatively transferred genes, we used four strict criteria as follows: 1) the gene must be absent in the control rhizospheres, 2) the gene must be detected in the inoculated rhizospheres, 3) the gene must be present in the inoculum and 4) the gene must be at least 128 times more abundant than it would have been expected for relic DNA from the inoculum. For this latter criterium, we calculated that the inoculated DNA (∼850 µg) would have been diluted in the DNA from the entire pot (average of 50 ng/µl per extract x 100 µl of DNA extract x 500 mg of soil per extraction x 1.5 kg of soil per pot = 15,000 µg of DNA per pot), for a 5.67% dilution factor. We then added a 128x safety margin (corresponding to 7 doubling events) and only considered genes that had a relative abundance in the inoculated rhizospheres that was at least 7.26 times (5.67% x 128) their relative abundance in the inoculum.

## Results

### Plant parameters

Plants growing under 15% of the SWHC (LW) had lower plant fresh biomass (F=15.9, P=0.00052), leaf water content (F=12.4, P=0.0017), and plant water content (F=26.8, P<0.001), as compared to plants growing under 50% of the SWHC (HW) (Fig. 1). The interaction term SWC:Inoculum was only significant for the super oxide dismutase (SOD) activity (F=4.0, P=0.032), with the differences being driven by the lower SOD activity in the LW plants as compared to the HW plants for the continuous inoculum and the inverse trend for the intermittent inoculum and the control (Fig. 1). Inoculation alone did not affect any of the plant parameters.

**Figure 1.**
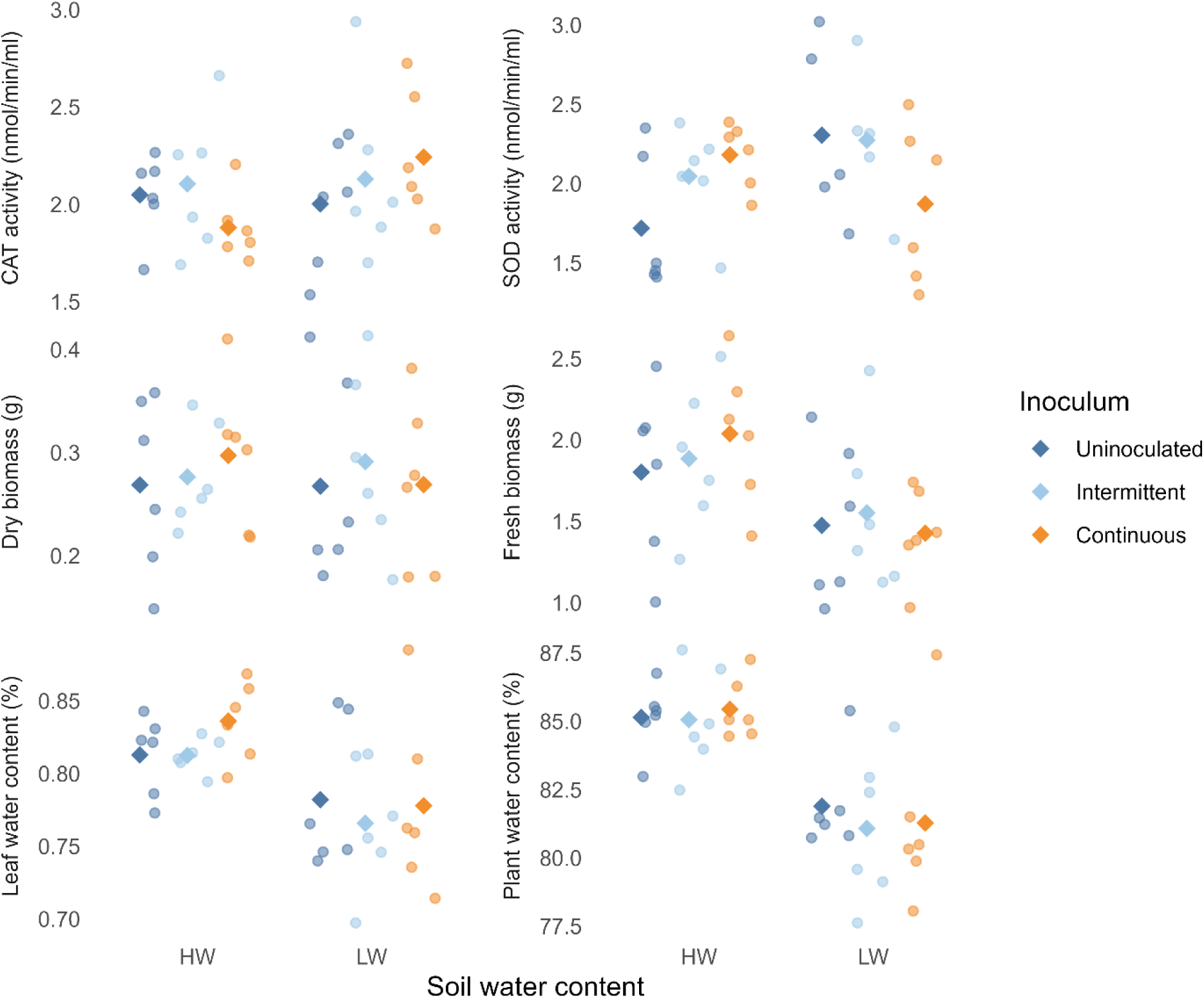
Plant parameters mostly respond to soil water content, with some effect of DNA inoculation on stress enzyme activities. Enzymatic activities for catalase (CAT) and superoxide dismutase (SOD), and plant fresh and dry biomass, and leaf and total water content following five weeks at 15% of the soil water holding capacity (SWHC) (LW) or 50% of the SWHC (HW), and three weeks after inoculation with DNA extracted from soil with a continuous or intermittent water stress history, or with water (uninoculated controls). Diamonds represent the arithmetic mean (n=6).

### Effect of the DNA inoculation on the microbial community

The main factor influencing the wheat rhizosphere microbial community was the soil water content, with a clear differentiation along the first axis of the PCoA ordinations (Fig. 2a). Permanova tests confirmed this visual interpretation (R^2^=0.099, P=0.001). The DNA inoculations did not change the wheat rhizosphere microbial community (R^2^=0.051, P=0.47). The community composition at the family level was very similar between inoculated and non-inoculated rhizosphere samples (20 most abundant families, Fig. 2b). Only one family showed a significant shift: the relative abundance of the *Hyphomicrobiaceae* (*Alphaproteobacteria*) increased to an average of 2.88% of the total community when inoculated with the continuous inoculum in the HW soils, as compared to an average of 2.14% in the controls (paired Wilcoxon test: W=21, P=0.031). Three other families showed nearly significant (0.05<P<0.10) shifts: in the HW soils, the continuous inoculum increased the relative abundance of *Mycobacteriaceae* (*Actinobacteria*, 6.02% vs. 5.70%, W=19, P=0.094) and decreased the relative abundance of the *Nocardiaceae* (*Actinobacteria*, 8.35% vs.9.57%, W=1, P=0.063), whereas in the 15% soil, both the continuous and the intermittent inoculums decreased the relative abundance of the *Nitrososphaeraceae* (*Thaumarcheota*, 7.20% vs. 7.49%, W=1, P=0.063, and 6.94% vs. 7.49%, W=2, P=0.094, respectively).

**Figure 2.**
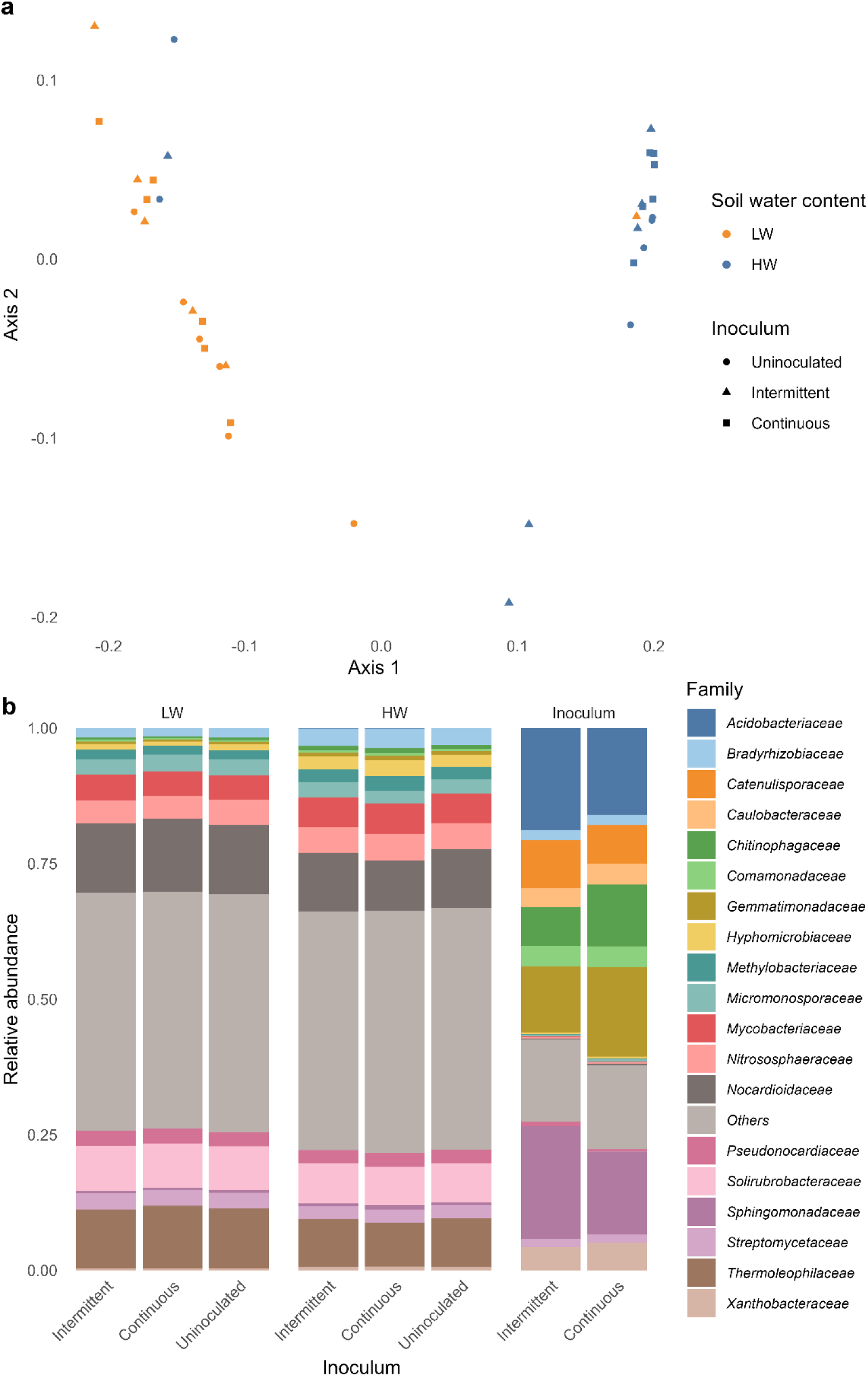
Rhizosphere microbial community of wheat plants are mostly affected by soil water content, with no effect of DNA inoculation. a) Principal coordinate analysis based on Bray-Curtis dissimilarity of the metagenomic-derived gene table and b) community composition for the top 20 most abundant families in the metagenomic-derived gene table, for rhizosphere samples of wheat growing under 15% of the soil water holding capacity (SWHC) (LW) or 50% of the SWHC (HW) after inoculation with DNA extracted from soil with a continuous or intermittent water stress history, or with water (uninoculated controls). In a), axis 1 and 2 represent 15.5% and 2.8% of the variation in the dataset, respectively.

### Laterally transferred genes in the wheat rhizosphere

We used WAAFLE [37] to identify potentially laterally transferred genes among the contigs created from the assembly of all samples, excluding the inoculum. We found 111 contigs that had clear signs of LGT, onto which 114 genes were potentially transferred (Supplementary Table S1). These laterally transferred genes were mostly from the *Actinobacteria* (81 genes, 71.1%) and *Proteobacteria* (24 genes, 21.1%), with one *Chloroflexi* and one *Gemmatimonadetes* gene, with the remaining (7 genes) being unassigned at this taxonomical level. These genes had a wide variety of KEGG definition, and most were unique to a single gene. Among the few functions that were found more than once at the KEGG pathway level, we found 8 genes linked to sulfur (biotin metabolism, cysteine and methionine metabolism, glutathione metabolism, sulfur metabolism, sulfur relay system, taurine and hypotaurine metabolism), 7 to amino acid metabolism, 6 to transporters, 5 to antimicrobials (vancomycin resistance, monobactam biosynthesis, biosynthesis of ansamycins, antimicrobial resistance genes, biosynthesis of type II polyketide products), 4 to butanoate metabolism, 4 to chaperones and folding catalysts, 3 to biofilms and quorum sensing, and 3 to transposases. None of the genes identified by WAAFLE had significant BLAST matches to genes from the inoculum assembly, suggesting that these genes were not recent transfers resulting from the inoculation, but rather LGT events from the receiving soil that predated our experiment. Accordingly, only gene_id_1256981, coding for a chaperonin (GroES), was significantly more abundant in the “intermittent” rhizosphere as compared to the controls (Padj=0.048 in paired Wilcoxon tests). The low soil water content treatment positively affected the relative abundance of 14 LGT genes (Supplementary Table S2), suggesting that they might have an adaptative value when water is scarce. Another 14 LGT genes were less abundant in the LW soils (Supplementary Table S2).

### Genes from the inoculum overrepresented in the wheat rhizosphere

We took a different approach to identify genes that would have been potentially integrated from the inoculum and might have been missed by WAAFLE because of their low abundance, low prevalence and/or presence in short contigs. We first re-mapped the reads of the rhizosphere metagenome on contigs created from the assembly of the inoculum metagenome alone. This dataset therefore only contained genes that were found in the inoculums. We then looked at genes that were absent from the control (uninoculated) rhizosphere and were present in the inoculated rhizosphere above a certain relative abundance, to make sure it was not DNA remnants from the inoculation (see material and methods for details). We did this separately for each sample, because transformation events would be expected to happen randomly across replicates. Using this method, we found 278 (average per sample of 27.6) and 258 (average per sample of 30.2) genes putatively transferred to the rhizosphere communities from the continuous and intermittent inoculums, respectively. This represents a transformation rate around 10^-11^ transformation events per cell (assuming 5 Mbp genomes each weighing 5 x 10^-6^ ng, which would mean 3 x 10^12^ microbial cells per pot). Most of the genes were flagged for only one treatment (236 for continuous and 182 for intermittent). Only 32 of the putatively transferred genes were defined at the KEGG entry level (Supplementary Table S2).

We compared the taxonomic affiliation of the transferred genes against the taxonomic make-up of the rhizosphere. We did this comparison sample per sample, pairing the putatively transferred genes from one sample to its corresponding rhizosphere sample. Among the putatively transferred genes, there was an overrepresentation of genes affiliated to the *Acidobacteria* (40.8 of the transferred genes vs. 8.6% of the rhizosphere reads, P=1.19 x 10^-7^), Bacteroidetes (9.14 vs. 0.44%, P=0.0018), and *Gemmatimondetes* (7.5 vs. 1.2%, P=0.032), and an underrepresentation of the *Actinobacteria* (5.4 vs. 54.4%, P=1.19 x 10^-7^), *Chloroflexi* (0.5% vs. 6.7%, P=1.19 x 10^-7^), and *Thaumarchaeota* (1.1 vs 4.8%, P=1.19 x 10^-6^) (Fig. 3a).

**Figure 3.**
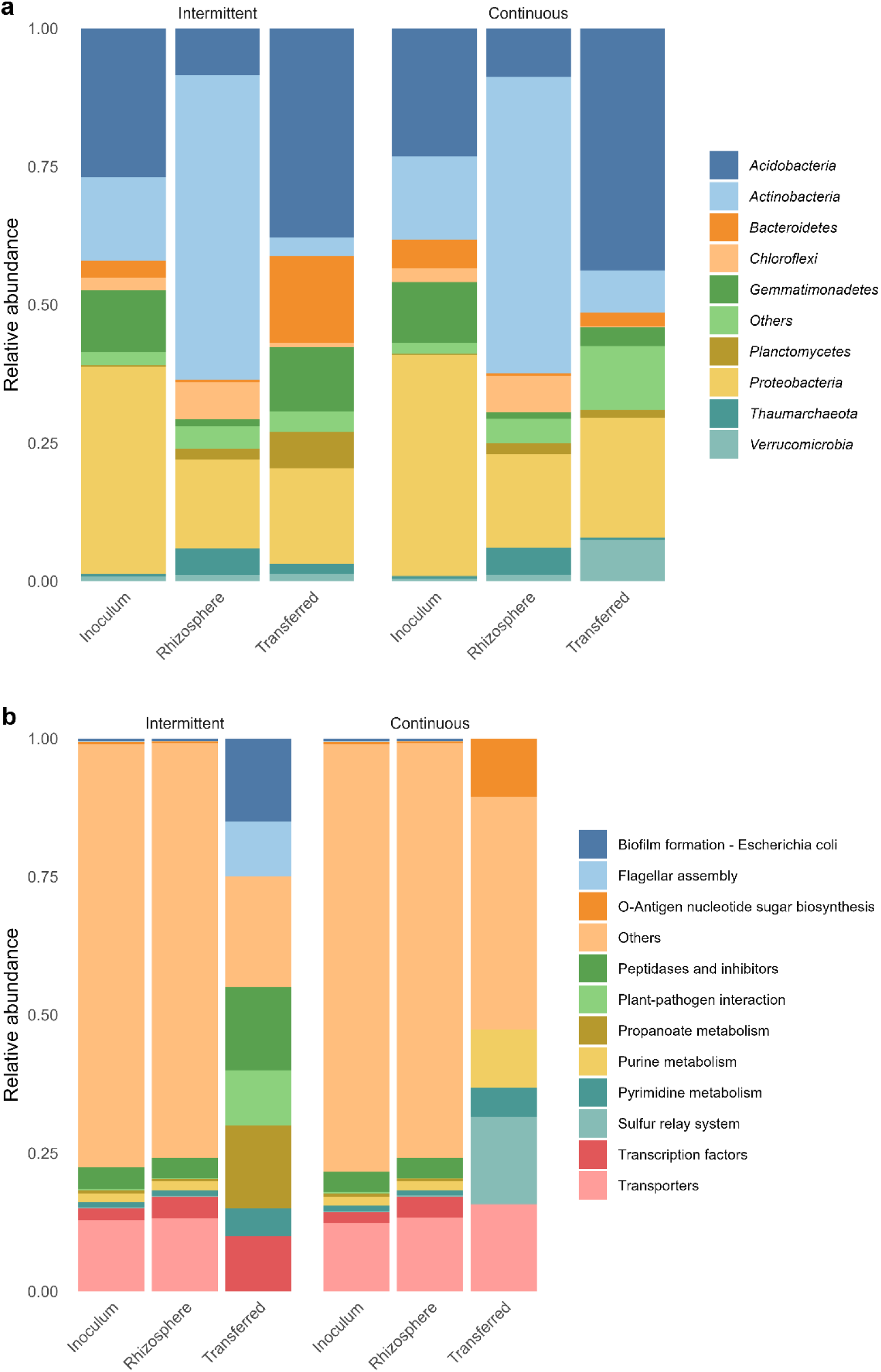
The putatively transferred genes are not a random subset of the taxa or function present in the rhizosphere or the inoculum. Community composition at the phylum (a) and KEGG entry (b) levels for the inoculum, the wheat rhizosphere and the putatively transferred genes, for the intermittent and continuous DNA inoculums.

Since very few of the putatively transferred genes could be annotated, we could not perform the same statistical analysis as for taxa. The KEGG pathways of the putatively transferred genes are shown in Fig. 3b alongside the affiliation of the all the genes in the rhizosphere and inoculum. Pathways related to propanoate, plant-pathogen interactions, sulfur, biofilm formation, transporters, peptidase and nucleic acid metabolism appeared to be overrepresented among the putatively transferred genes.

There was no gene that was identified by both approaches (WAAFLE and abundance analysis), but there was an overlap between the datasets at the level of two KEGG entries: chaperonin GroES, and the meso-butanediol dehydrogenase / (S,S)-butanediol dehydrogenase / diacetyl reductase, from the butanoate metabolism.

## Discussion

As in Giard-Laliberté [23], we compared the capacity of the microbial communities of two highly similar wheat field soils with contrasting water stress history [31, 32, 38] to rescue wheat plants growing under water stress. However, instead of extracting and inoculating live microbes, we extracted and inoculated large quantities of DNA, with the idea that the rhizosphere microbial community would take up genetic elements from this inoculated DNA. Natural genetic transformation in soils has mostly been studied to determine if genes from genetically modified organisms (GMOs) could be transferred to the soil microbial communities [39]. Here, even though the effects were subtle, we showed convincing evidence of natural genetic transformation from the inoculated DNA to the wheat rhizosphere microbial communities. Indeed: 1) several hundreds of genes from the inocula were exclusively present in the rhizosphere of the inoculated plants, at levels well above what could be expected for relics of the inoculated DNA, 2) many of the genes that were identified as putatively transferred and could be assigned to a function were not a random subset of the whole community but rather linked to life in the plant/soil environment, and 3) one of the inoculum reduced plant stress levels under low soil water content.

One of our concerns with the current experiment was about DNA persistence in soil. If DNA could not survive long enough in the soil, the microbial communities would not have been able to take it up. Alternatively, if the half life of DNA in soil was too long, the transformation events detected could have been due to the sequencing of residual DNA from the inoculum. The first possibility is not supported by the literature, as DNA appears to maintain itself in soil for relatively long periods of time, with the presence of transgenic DNA from plant material being detected 30 days after incorporation of leaf biomass to soil microcosms [40], and up to 1 year in soil where transgenic crops were cultivated [41]. Ultimately, DNA half-life is influenced by the soil physico-chemical properties and the microbial communities [42], but the apparent uptake of inoculated DNA by the microbial community indicates that the inoculated DNA survived long enough in the soil. The alternate possibility that the inoculated DNA would persist in the soil and be detected and mistaken for transformation events in the rhizosphere was controlled by only considering the genes that were more than 7 times above the expected abundance for inoculated DNA that would have simply persisted in the soil. Another, indirect indication that the genes reported were not from the inoculated DNA that had lingered in the soil, is the non-random assortment of functions, with most of the genes being linked to functioning in the plant/soil environment, in agreement with our general hypothesis.

Some of the genes identified as putatively transferred were related to defense systems or antimicrobial resistance. The soils used to create the inoculant were from a semi-arid region and were either constantly or intermittently exposed to dry conditions. Soils under drought were previously reported to contain more antibiotics [43]. This was hypothesized to be linked with a drought-induced increase in the relative abundance of recognized antibiotic producers such as the *Actinobacteria*, which also overproduce antibiotics when faced with osmotic stress [44, 45].

Under limited resource availability such as in the soil, highly competitive organisms producing antibiotics, have a clear advantage [46], so it would make sense for such genes to be taken up and integrated. Even under water-replete conditions, the uptake of antibiotic production genes is advantageous and would increase the competitiveness of the transformants. Other putatively transferred genes were related to the production of extracellular polysaccharides and to biofilm formation. Bacteria form biofilm to cope with environmental stresses, but it is also an important adaptation for the life in the root environment [47]. A gene from the starch utilization system (SUS) that includes proteins for binding and degrading starch at the surface of the cell were also found among the putatively transferred genes. Starch is a common polysaccharide in plant residue and is also is one of the components of root mucilage, which increases rhizosphere water content and plays a key role in the adaptation of plants to drought [48, 49]. Some other genes singled out as potentially transferred were linked to the production of osmolytes, such as genes linked to betaine biosynthesis or amino acid metabolism. Genes related to the metabolism of carboxylic acids – key root exudates [50] – were also putatively transferred. More specifically, we found genes related to the propanoate and butanoate metabolism, and, in the case of the butanoate, we found a key gene (K18009-budC) coding for a protein that can reduce diacetyl – a volatile molecule that was shown to increase following drought in soils [51] – and can produce butanediol or acetoin – which can either confer drought resistance to plants [52] or promote plant growth [53]. A gene coding for a chaperone was also identified, which have important roles in the response to drought stress for both microbes and plants [54–56].

We also used WAAFLE, a tool designed to find LGT genes among contigs and found several genes that were putatively transferred. However, none of these genes were found in the inoculum, suggesting they were older LGT events that happened in the recipient soil, before the inoculation. Since the recipient soil was never subjected to drought, we consider the genes found by WAAFLE as having an importance for life in the soil environment. Although none of the genes were identical, two similar KEGG entries, among the most interesting genes discussed above, were found by the methods: *budC* and a chaperon. One of the *budC* gene identified by WAAFLE was relatively more abundant in the 15% SWHC treatment, suggesting that it might be important for adaptation to low water conditions. On top of these, there was also a large overlap at higher functional categories, with the two methods highlighting genes related to antimicrobials, amino acids, chaperones, butanoate metabolism, biofilms, and sulfur metabolism. This further confirms our hypothesis and strengthen our results: the genes that we think were taken up from the inoculum are functionally similar to the ones that are transferred and get fixed in a soil community in the longer term. One key difference was, however, the taxonomic affiliation of the transferred genes between the two approaches. For WAAFLE, most of the transferred genes were found in *Actinobacteria* and *Proteobacteria* contigs, whereas using our abundance analyses, most of the putatively transferred genes were linked to the *Acidobacteria* and *Bacteroidetes*. This might highlight different ecological strategies of major soil phyla.

We previously showed that inoculated bacteria that persisted in the plant environment under drought stress had larger genomes with more genes, as compared to bacteria that did not persist [16]. Bacteria and archaea can adapt to changing environmental conditions by having two or more copies of a gene, dubbed ecoparalogs, each encoding for a protein having a different environmental optimum [57]. Ecoparalogs could arise from the duplication and divergence of an ancestral gene but could also arise from LGT [57]. This accumulation of ecoparalogs seem to have been the evolutionary strategy behind the success in terrestrial environments of some bacterial groups, such as the *Acidobacteria*. Indeed, genomes of acidobacterial strains isolated from soils typically harbour a larger genome size and a higher proportion of paralogous genes compared to strains from other environments [58, 59]. Additionally, genome analyses have shown a high level of LGT among *Acidobacteria* [58], shaping the genomes and leading to the acquisition of new genetic potential (such as auxiliary metabolic genes) [60]. Our results also showed an overrepresentation of *Acidobacteria* in the genes flagged as putatively transferred from our inoculum. On average, only 8.6% of the genes could be attributed to this phylum in our wheat rhizospheres, as compared to up to 40% of the potentially transferred genes. The major phyla, such as *Actinobacteria* and *Proteobacteria* were not well represented among the genes potentially transferred from the inoculum, even though they dominated the rhizosphere. This underscores the fact that the uptake of genes from the environment might be a strategy used by some defined taxonomic groups, which would give them an advantage during resource fluctuations in soils. Here, we provide preliminary evidence that natural genetic transformation in soil *Acidobacteria* is not only part of their evolutionary history, but that it might also be a contemporary mechanism these bacteria have evolved to adapt rapidly to changing environments. The fact that 1) transformation events mostly occurred in underrepresented phyla, 2) *Acidobacteria* are typically decreasing in relative abundance with increasing soil water stress [32], and 3) *Acidobacteria* are not generally recognized as being able to help plant withstand water stress might partly explain why we did not find significant effects of the inoculations on the community structure, and little effects on the plants.

The only effect on plants was a reduced level of superoxide dismutase – suggesting less water stress – for the “continuous” inoculum under low soil water content. In other experiments using the same soils used here to create the inoculums, the rhizosphere of plants growing in the “continuous” soil was enriched in genes related to antioxidants, as compared to the “intermittent” rhizospheres [61]. Similarly, the decrease in leaf water content following reduced soil water content was less important for wheat grown in the “continuous” field, as compared to the “intermittent” field [38]. The microbial communities of the two soils also responded differently to a subsequent exposure to water stress, both in terms of activity and community composition [31, 32]. We had also previously showed that under water stress, most of the transcriptomic response in the root-rhizosphere environment was due to bacteria, with many osmolyte-related genes being differentially expressed [2]. We had also showed that inoculating highly osmotolerant bacteria allowed wheat plants to conserve more water in their tissues when faced with drought [16]. There is some evidence that microbial endophytes and rhizobacteria can increase plant osmolyte concentration [62, 63], including proline [64], and some studies have reported that microbes can exude these compounds in the plant environment [65, 66], enabling them to directly contribute to the plant osmolyte concentration during water stress. Although there are some prime suspects, more work will be needed to pinpoint which of the potentially transferred gene could have reduced plant stress levels.

For other plant parameters, the transformation events did not lead to clear effects on the plants. A first possible cause was that transformation events were rare and only a small proportion of the microbial community was transformed (we calculated a rate of 10^-11^ successful transformation per cell), in line with previous reports that evaluated the occurrence of naturally transformable bacteria in the rhizosphere to be negligibly low [29] or relatively rare [30]. As discussed above, this could be linked to different ecological/evolutionary strategies across phyla, but one could also speculate that natural genetic transformation is performed by populations under a certain density to tip the competitive balance to their profit. Additionally, we had hypothesized that the rhizosphere microbial communities would be more prone to be transformed naturally when under water stress. Transferred genes can become rapidly fixed in a population under selective conditions, even if LGT events are rare, as it is the case for the rapid spread of antibiotic resistance following exposure of microbial populations to antibiotics and of xenobiotics degradation in polluted soils [67]. Although there were no differences in terms of quantity, we did find differences in the KEGG classification of the potentially transferred genes between the low and high soil water treatments, and, as mentioned above, the reduction in plant SOD activity was only observed under low soil water content, partly confirming this hypothesis.

## Conclusion

Our results are pointing out that natural genetic transformation might occur between well established microbial communities and exogenously applied DNA. The genes putatively transferred from the inoculum were not a random assortment of genes, but rather had functions linked to microbial life in soil and were mostly found in a key soil phylum, the *Acidobacteria*.

One of the inoculations resulted in a reduction of wheat stress under low soil water content, suggesting that the natural genetic transformation of established rhizosphere communities can have beneficial effects on the plant. This is an exciting first step toward harnessing LGT for microbial community modifications. With further refinements, this approach could be used to modulate crops’ phenotype, toward microbio-centric sustainable agriculture.

## Acknowledgements

We would like to thank all past and present members of the mECO:LABS for support and helpful discussions.

## Funding

This study was supported by a Natural Science and Engineering Research Council of Canada (NSERC) Discovery Grant (2014-05274), a NSERC Strategic grant for project (STPGP 494702) and a Compute Canada Resource allocation on the Graham system (2018-714) to E.Y. C.G.-L. was supported by a NSERC Alexander Graham Bell Canada Graduate Scholarship.

## Competing interests

The authors declare no competing financial interests in relation to the work described.

## Data availability

Raw sequence data are available in the Short Read Archive (SRA) of the NCBI under Bioproject PRJNA624060. The metagenome co-assemblies and intermediate data needed to run the R scripts are avaible in Zenodo: https://doi.org/10.5281/zenodo.3755973. The R scripts used to manipulate data, perform statistical analyses and generate tables and figures are available on GitHub: https://github.com/le-labo-yergeau/DNA_Inoculation.

---

**Table S1:**
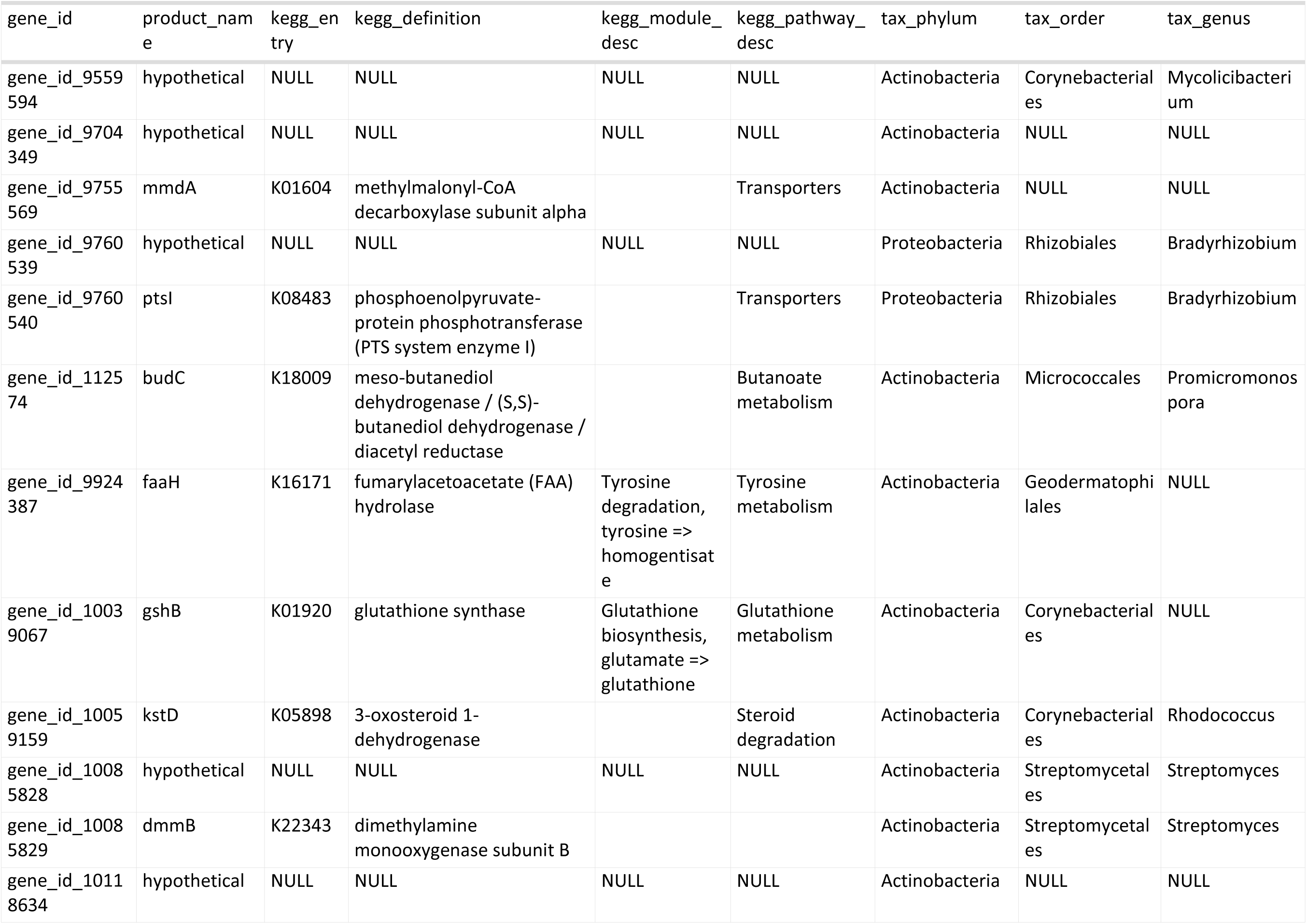
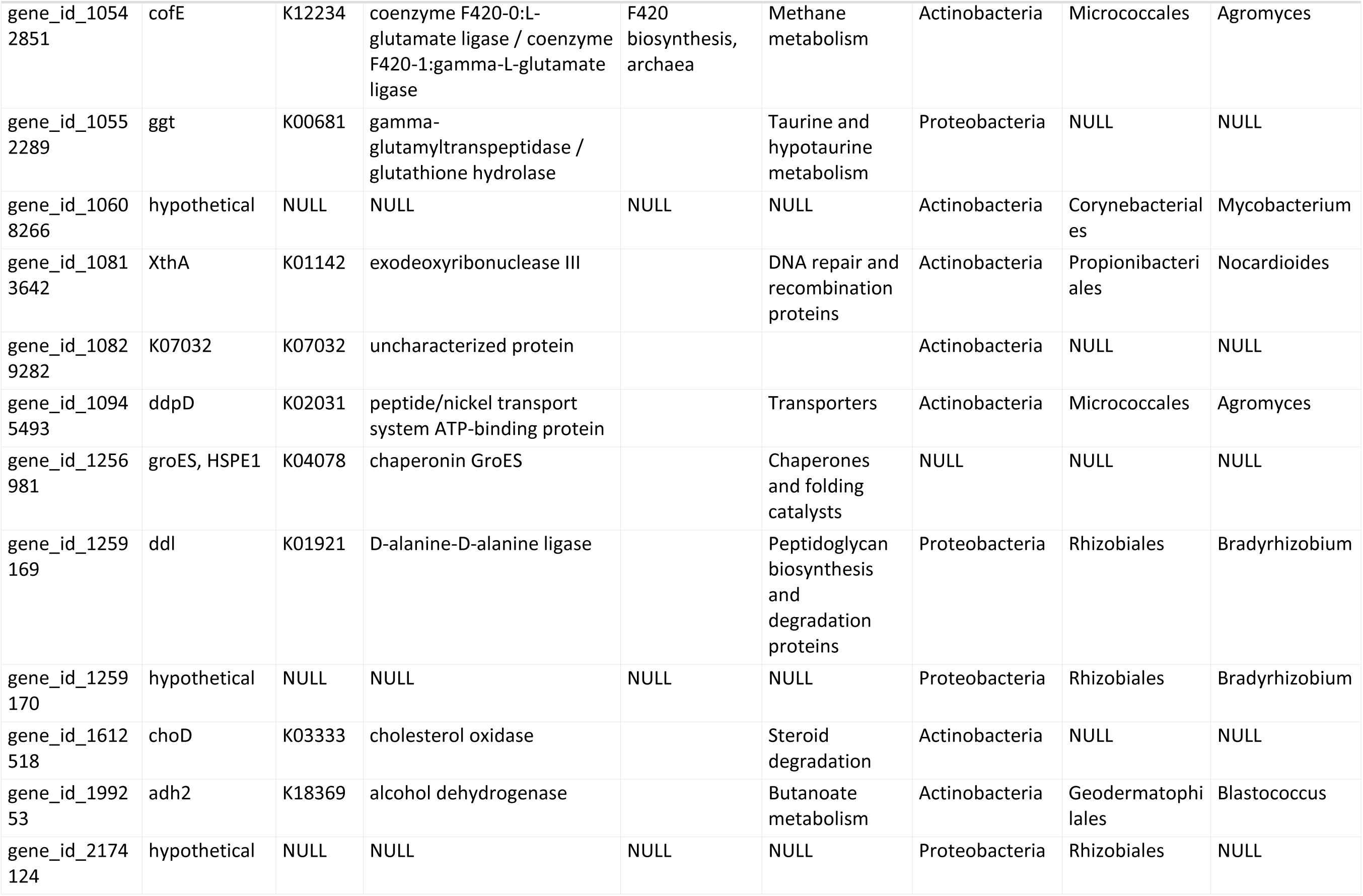
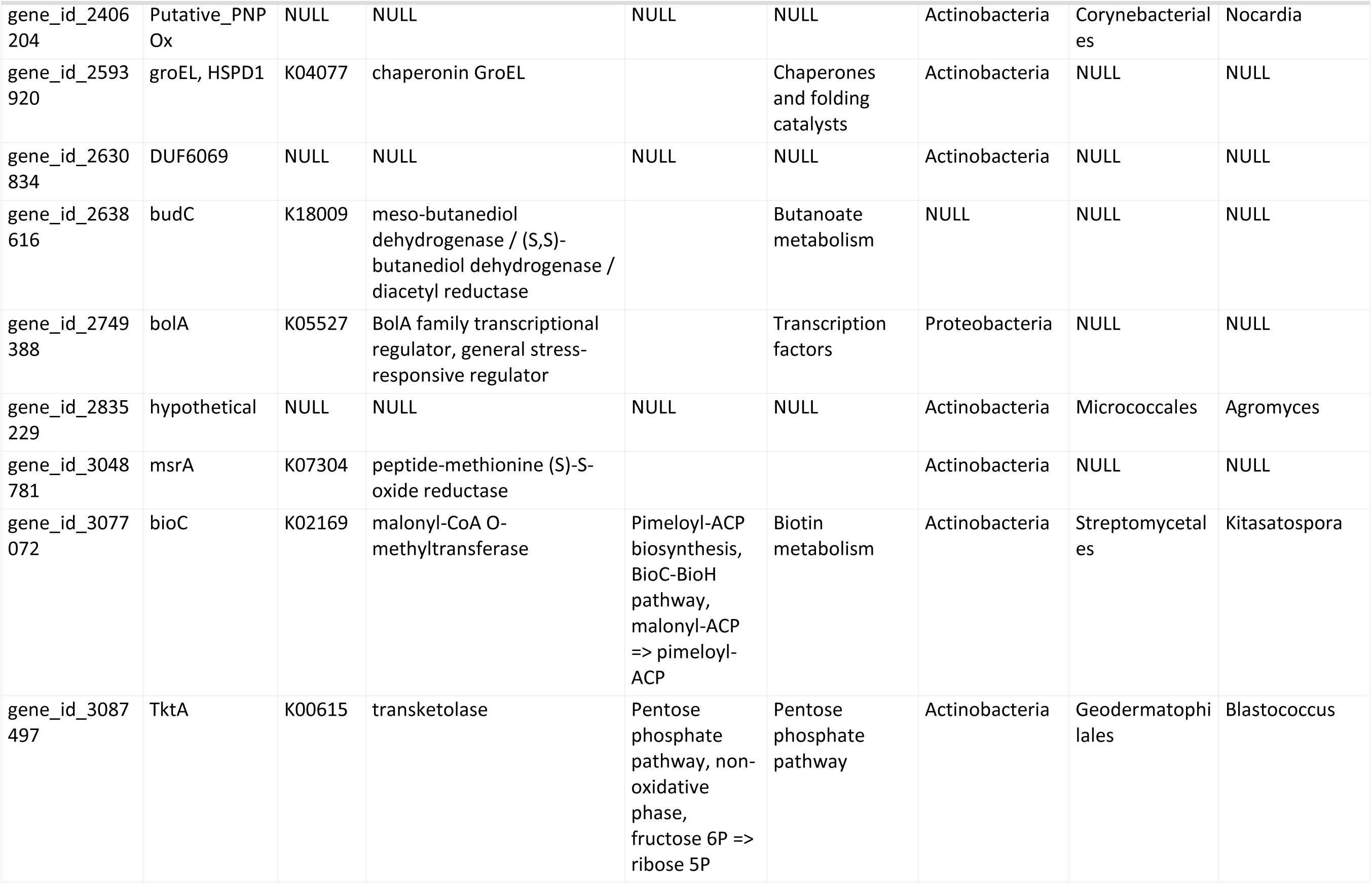
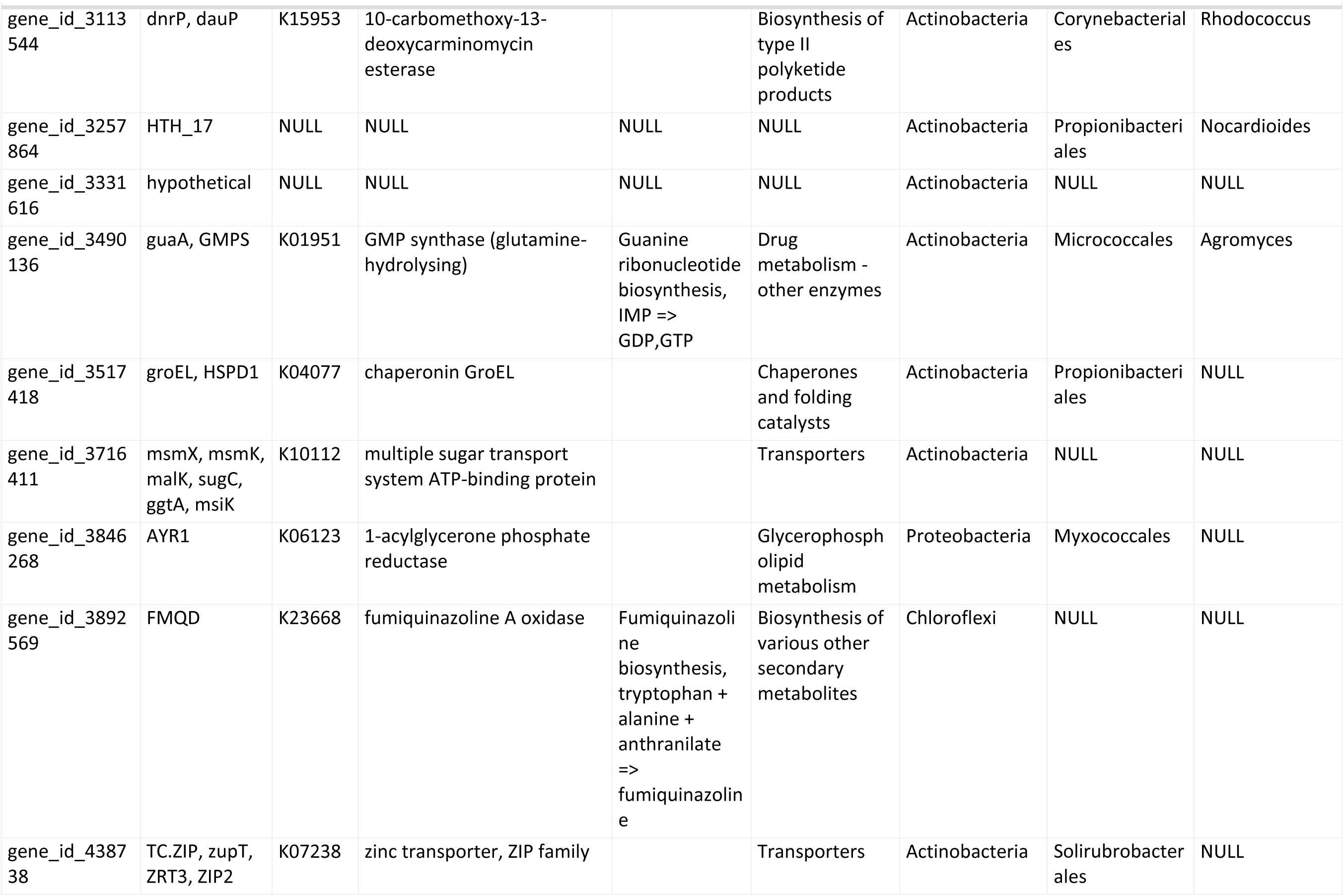
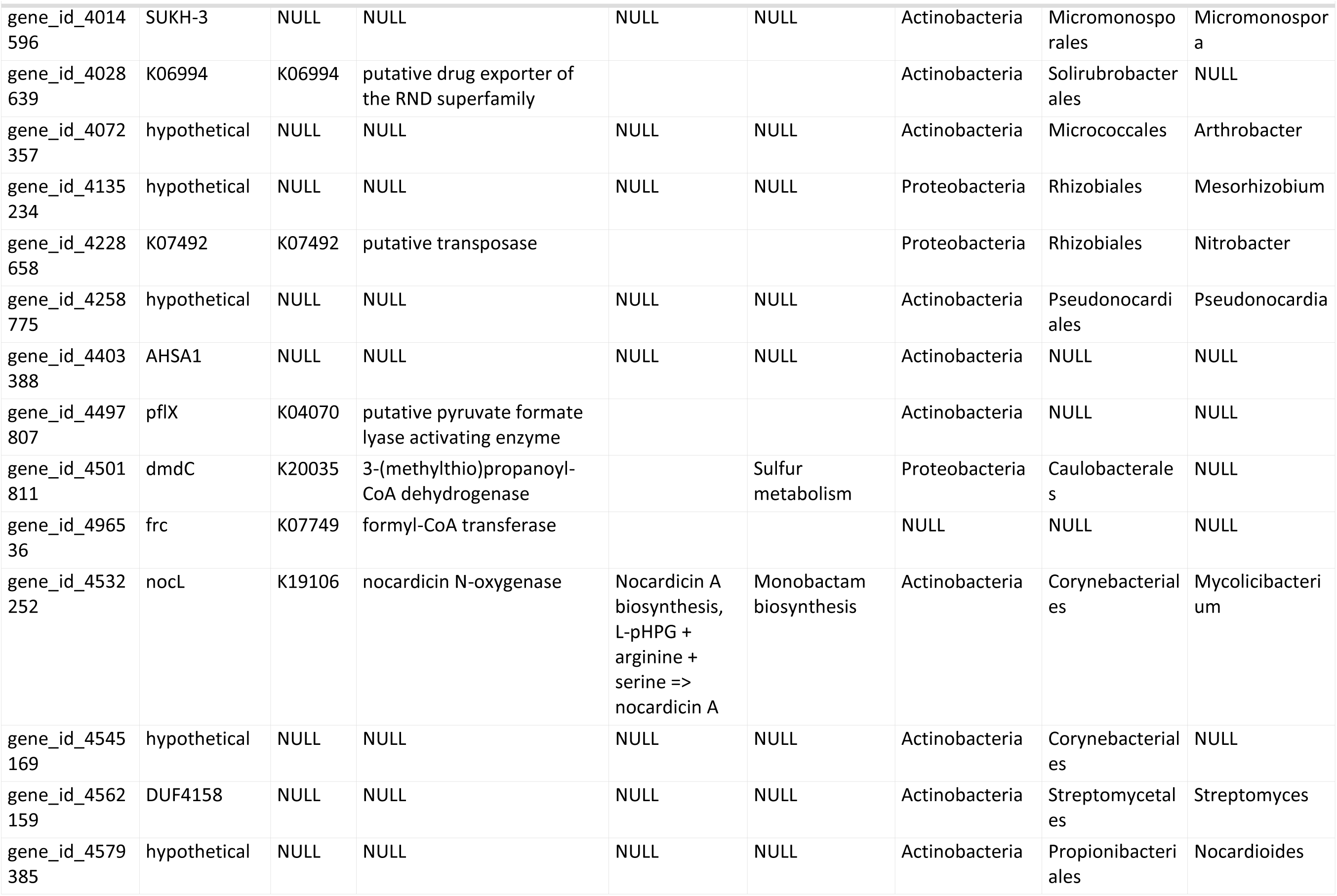
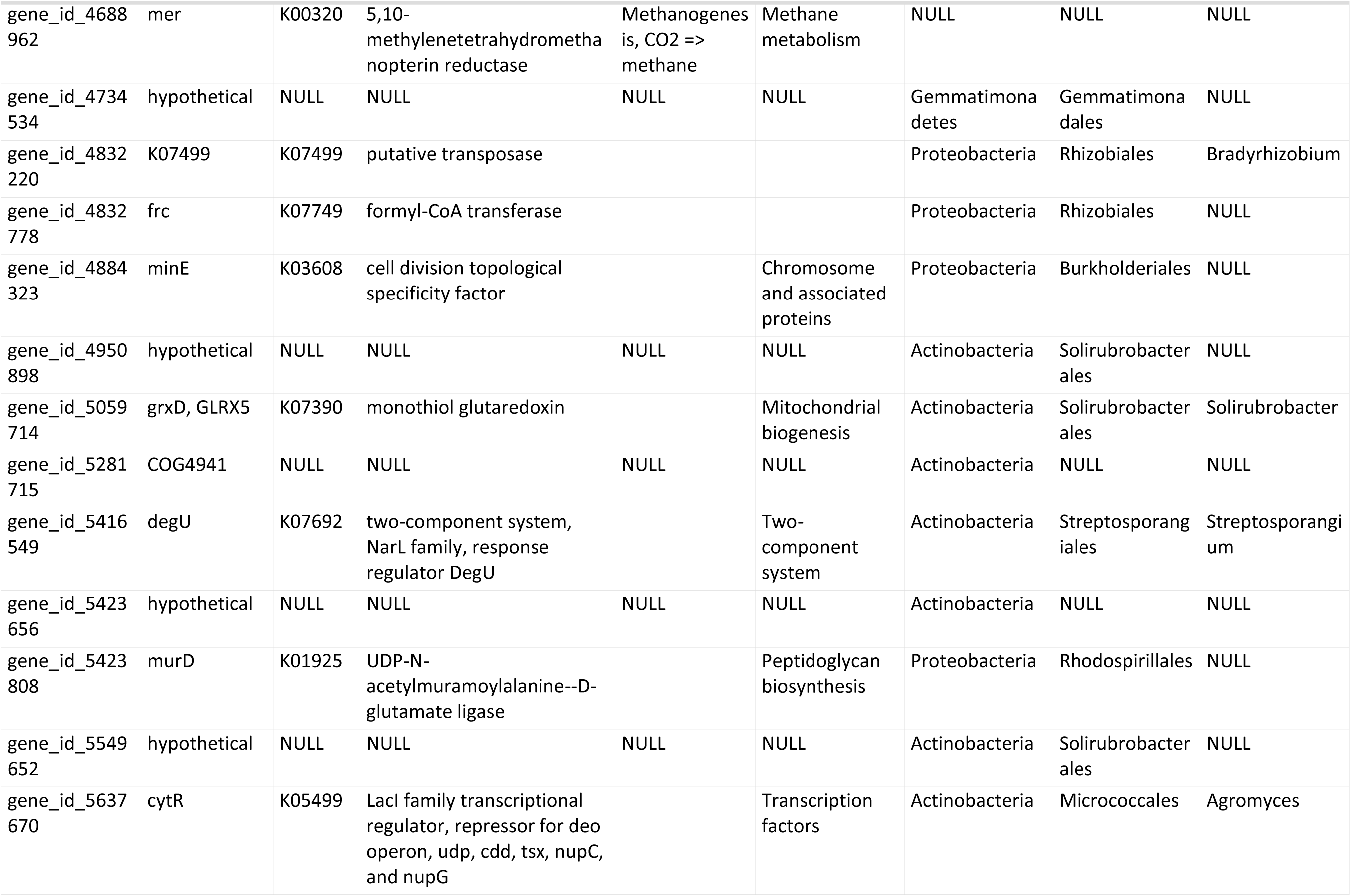
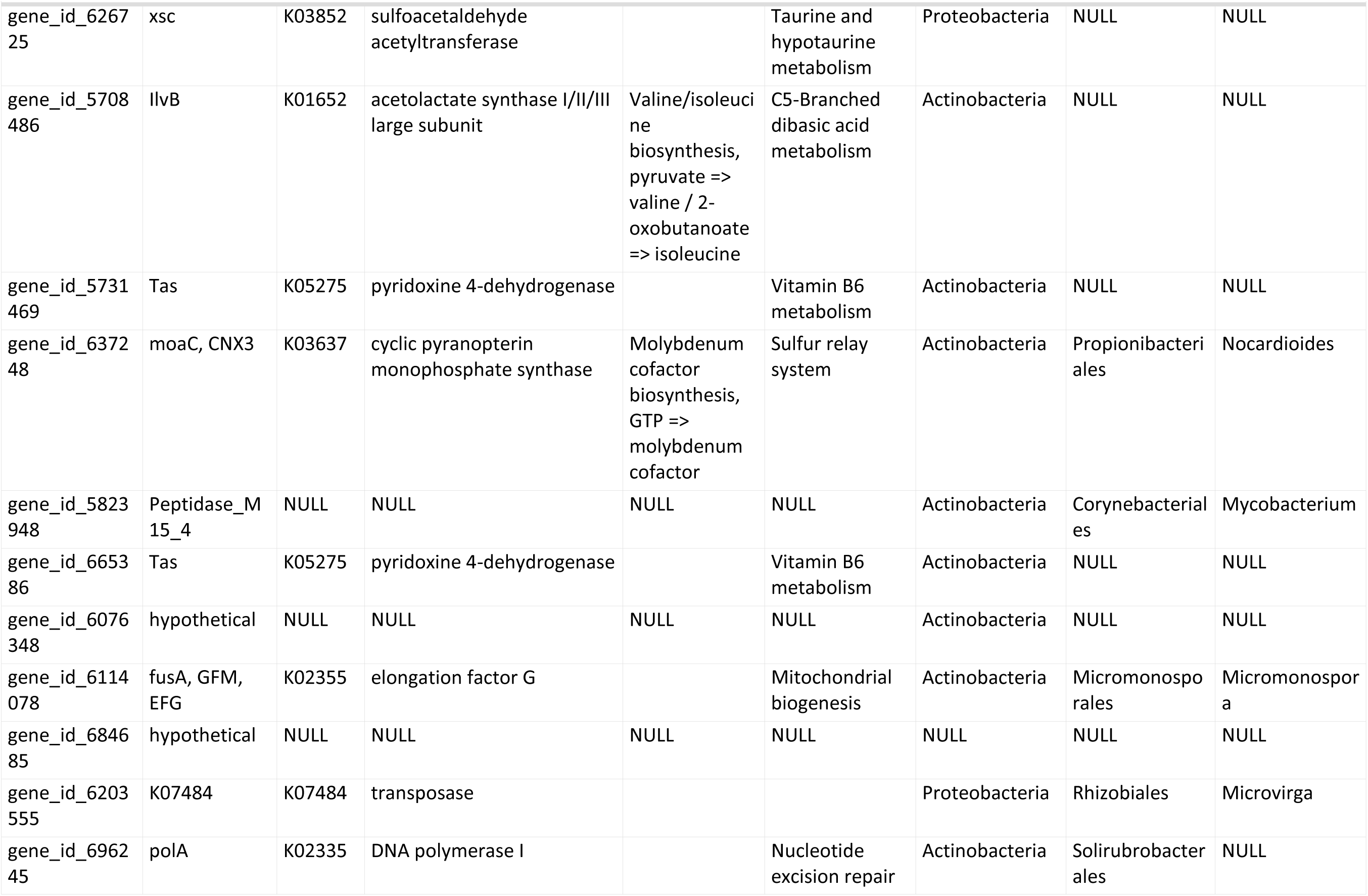
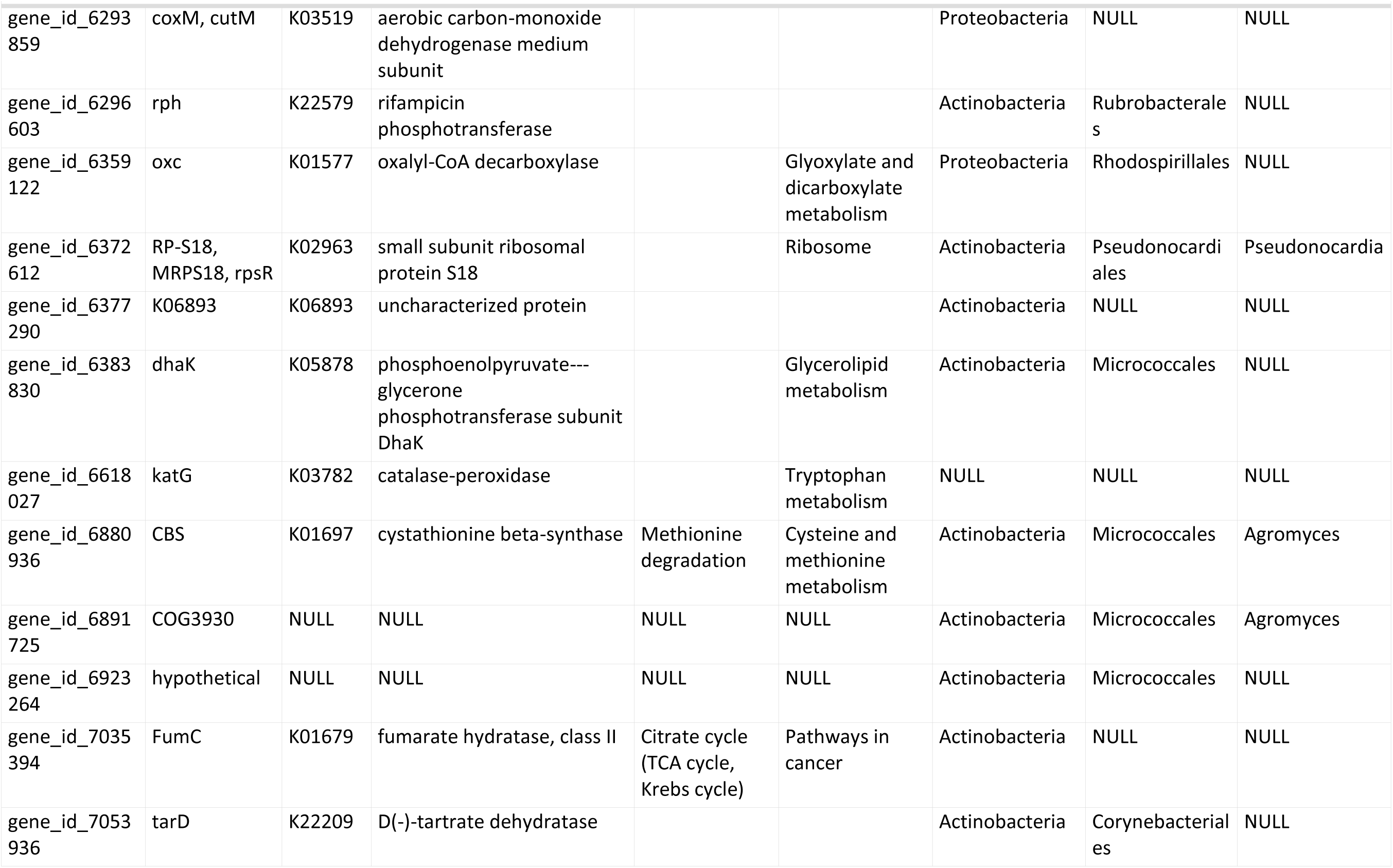
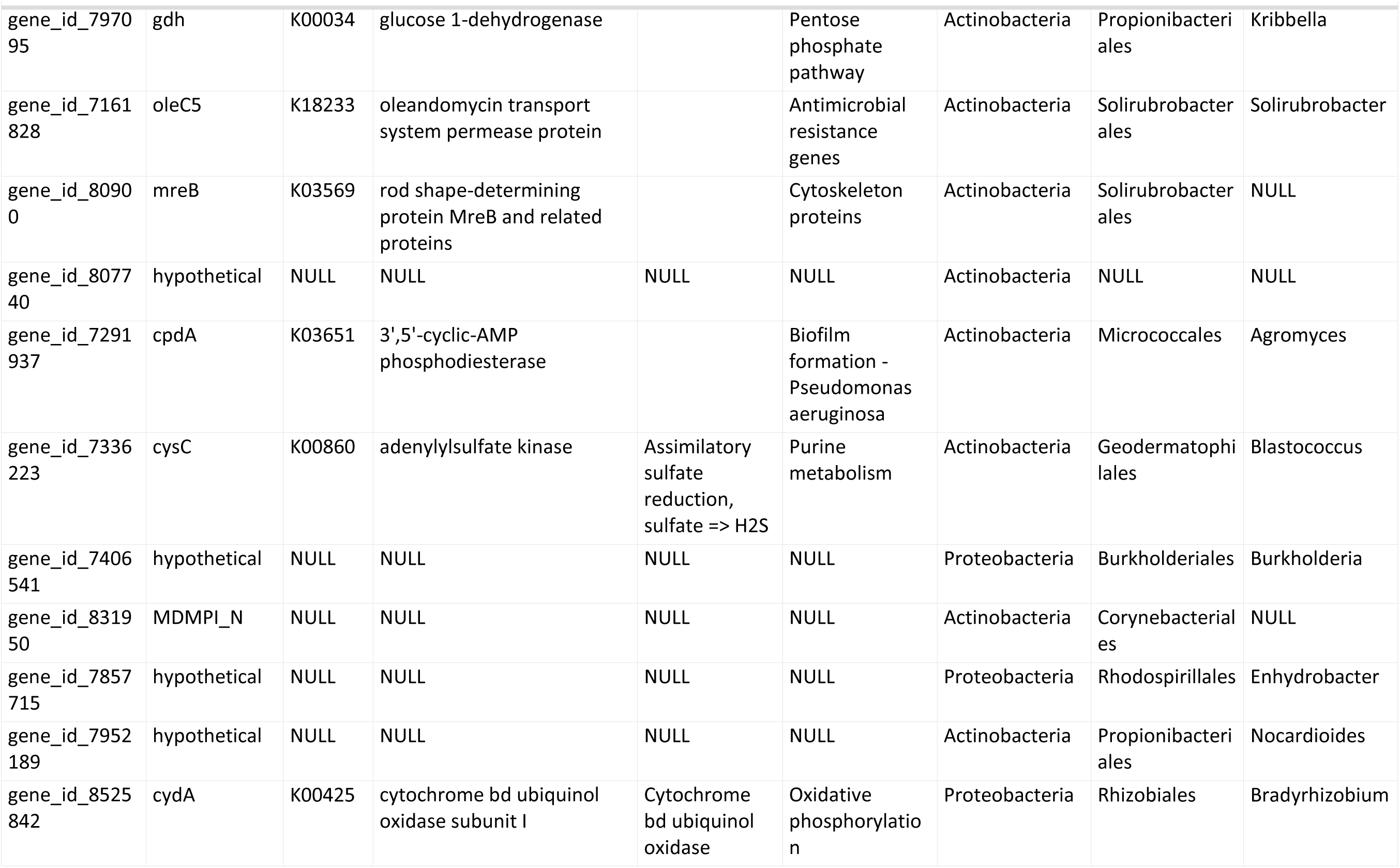
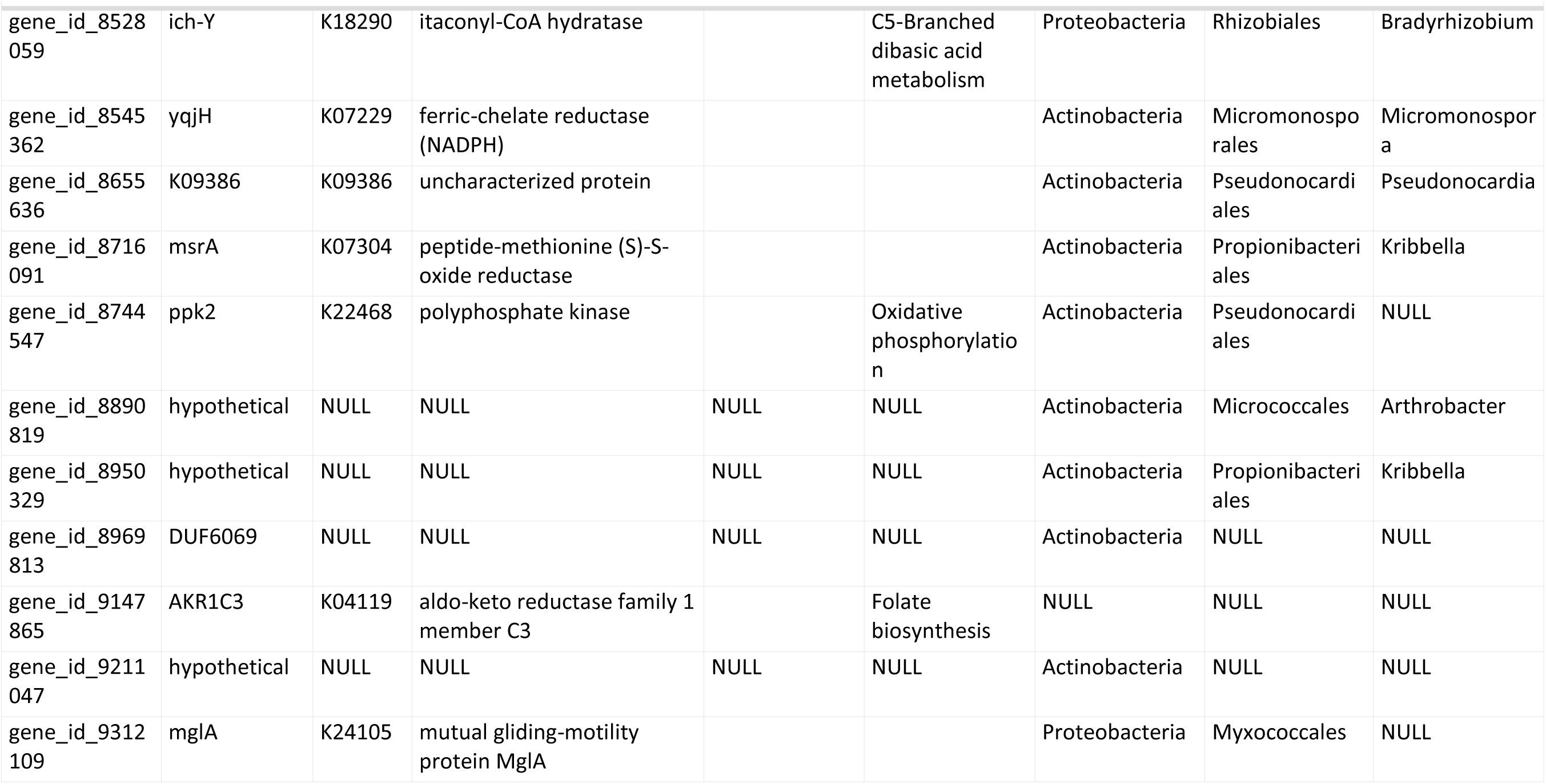
Potentially LGT genes identified by WAAFLE.

**Table S2:** Potentially LGT genes identified by WAAFLE that are significantly affected by soil water content.

| SWC | gene_id | n | mean | product_name | kegg_entry | kegg_definition | kegg_module_desc | kegg_pathway_desc | tax_phylum | tax_genus |
| --- | --- | --- | --- | --- | --- | --- | --- | --- | --- | --- |
| 15% | gene_id_10118634 | 18 | 0.063 | hypothetical | NULL | NULL | NULL | NULL | Actinobacteria | NULL |
| 50% | gene_id_10118634 | 18 | 0.041 | hypothetical | NULL | NULL | NULL | NULL | Actinobacteria | NULL |
| 15% | gene_id_112574 | 18 | 0.178 | budC | K18009 | meso-butanediol dehydrogenase / (S,S)-butanediol dehydrogenase / diacetyl reductase |  | Butanoate metabolism | Actinobacteria | Promicromonospora |
| 50% | gene_id_112574 | 18 | 0.078 | budC | K18009 | meso-butanediol dehydrogenase / (S,S)-butanediol dehydrogenase / diacetyl reductase |  | Butanoate metabolism | Actinobacteria | Promicromonospora |
| 15% | gene_id_1259169 | 18 | 0.194 | ddl | K01921 | D-alanine-D-alanine ligase |  | Peptidoglycan biosynthesis and degradation proteins | Proteobacteria | Bradyrhizobium |
| 50% | gene_id_1259169 | 18 | 0.363 | ddl | K01921 | D-alanine-D-alanine ligase |  | Peptidoglycan biosynthesis and degradation proteins | Proteobacteria | Bradyrhizobium |
| 15% | gene_id_1259170 | 18 | 0.165 | hypothetical | NULL | NULL | NULL | NULL | Proteobacteria | Bradyrhizobium |
| 50% | gene_id_1259170 | 18 | 0.362 | hypothetical | NULL | NULL | NULL | NULL | Proteobacteria | Bradyrhizobium |
| 15% | gene_id_199253 | 18 | 0.232 | adh2 | K18369 | alcohol dehydrogenase |  | Butanoate metabolism | Actinobacteria | Blastococcus |
| 50% | gene_id_199253 | 18 | 0.164 | adh2 | K18369 | alcohol dehydrogenase |  | Butanoate metabolism | Actinobacteria | Blastococcus |
| 15% | gene_id_2630834 | 18 | 1.008 | DUF6069 | NULL | NULL | NULL | NULL | Actinobacteria | NULL |
| 50% | gene_id_2630834 | 18 | 0.728 | DUF6069 | NULL | NULL | NULL | NULL | Actinobacteria | NULL |
| 15% | gene_id_2749388 | 18 | 0.039 | bolA | K05527 | BolA family transcriptional regulator, general stress-responsive regulator |  | Transcription factors | Proteobacteria | NULL |
| 50% | gene_id_2749388 | 18 | 0.119 | bolA | K05527 | BolA family transcriptional regulator, general stress-responsive regulator |  | Transcription factors | Proteobacteria | NULL |
| 15% | gene_id_3077072 | 18 | 0.127 | bioC | K02169 | malonyl-CoA O-methyltransferase | Pimeloyl-ACP biosynthesis, BioC-BioH pathway, malonyl-ACP => pimeloyl-ACP | Biotin metabolism | Actinobacteria | Kitasatospora |
| 50% | gene_id_3077072 | 18 | 0.090 | bioC | K02169 | malonyl-CoA O-methyltransferase | Pimeloyl-ACP biosynthesis, BioC-BioH pathway, malonyl-ACP => pimeloyl-ACP | Biotin metabolism | Actinobacteria | Kitasatospora |
| 15% | gene_id_3846268 | 18 | 0.005 | AYR1 | K06123 | 1-acylglycerone phosphate reductase |  | Glycerophospholipid metabolism | Proteobacteria | NULL |
| 50% | gene_id_3846268 | 18 | 0.014 | AYR1 | K06123 | 1-acylglycerone phosphate reductase |  | Glycerophospholipid metabolism | Proteobacteria | NULL |
| 15% | gene_id_3892569 | 18 | 0.046 | FMQD | K23668 | fumiquinazoline A oxidase | Fumiquinazoline biosynthesis, tryptophan + alanine + anthranilate => fumiquinazoline | Biosynthesis of various other secondary metabolites | Chloroflexi | NULL |
| 50% | gene_id_3892569 | 18 | 0.022 | FMQD | K23668 | fumiquinazoline A oxidase | Fumiquinazoline biosynthesis, tryptophan + alanine + anthranilate => fumiquinazoline | Biosynthesis of various other secondary metabolites | Chloroflexi | NULL |
| 15% | gene_id_4014596 | 18 | 0.110 | SUKH-3 | NULL | NULL | NULL | NULL | Actinobacteria | Micromonospora |
| 50% | gene_id_4014596 | 18 | 0.061 | SUKH-3 | NULL | NULL | NULL | NULL | Actinobacteria | Micromonospora |
| 15% | gene_id_4228658 | 18 | 0.199 | K07492 | K07492 | putative transposase |  |  | Proteobacteria | Nitrobacter |
| 50% | gene_id_4228658 | 18 | 0.333 | K07492 | K07492 | putative transposase |  |  | Proteobacteria | Nitrobacter |
| 15% | gene_id_438738 | 18 | 0.222 | TC.ZIP, zupT, ZRT3, ZIP2 | K07238 | zinc transporter, ZIP family |  | Transporters | Actinobacteria | NULL |
| 50% | gene_id_438738 | 18 | 0.139 | TC.ZIP, zupT, ZRT3, ZIP2 | K07238 | zinc transporter, ZIP family |  | Transporters | Actinobacteria | NULL |
| 15% | gene_id_4403388 | 18 | 0.240 | AHSA1 | NULL | NULL | NULL | NULL | Actinobacteria | NULL |
| 50% | gene_id_4403388 | 18 | 0.182 | AHSA1 | NULL | NULL | NULL | NULL | Actinobacteria | NULL |
| 15% | gene_id_4688962 | 18 | 0.429 | mer | K00320 | 5,10-methylenetetrahydromethanopterin reductase | Methanogenesis, CO2 => methane | Methane metabolism | NULL | NULL |
| 50% | gene_id_4688962 | 18 | 0.300 | mer | K00320 | 5,10-methylenetetrahydromethanopterin reductase | Methanogenesis, CO2 => methane | Methane metabolism | NULL | NULL |
| 15% | gene_id_4832778 | 18 | 0.019 | frc | K07749 | formyl-CoA transferase |  |  | Proteobacteria | NULL |
| 50% | gene_id_4832778 | 18 | 0.051 | frc | K07749 | formyl-CoA transferase |  |  | Proteobacteria | NULL |
| 15% | gene_id_4950898 | 18 | 0.852 | hypothetical | NULL | NULL | NULL | NULL | Actinobacteria | NULL |
| 50% | gene_id_4950898 | 18 | 0.603 | hypothetical | NULL | NULL | NULL | NULL | Actinobacteria | NULL |
| 15% | gene_id_496536 | 18 | 0.061 | frc | K07749 | formyl-CoA transferase |  |  | NULL | NULL |
| 50% | gene_id_496536 | 18 | 0.206 | frc | K07749 | formyl-CoA transferase |  |  | NULL | NULL |
| 15% | gene_id_5423808 | 18 | 0.081 | murD | K01925 | UDP-N-acetylmuramoylalanine--D-glutamate ligase |  | Peptidoglycan biosynthesis | Proteobacteria | NULL |
| 50% | gene_id_5423808 | 18 | 0.182 | murD | K01925 | UDP-N-acetylmuramoylalanine--D-glutamate ligase |  | Peptidoglycan biosynthesis | Proteobacteria | NULL |
| 15% | gene_id_626725 | 18 | 0.203 | xsc | K03852 | sulfoacetaldehyde acetyltransferase |  | Taurine and hypotaurine metabolism | Proteobacteria | NULL |
| 50% | gene_id_626725 | 18 | 0.545 | xsc | K03852 | sulfoacetaldehyde acetyltransferase |  | Taurine and hypotaurine metabolism | Proteobacteria | NULL |
| 15% | gene_id_6359122 | 18 | 0.148 | oxc | K01577 | oxalyl-CoA decarboxylase |  | Glyoxylate and dicarboxylate metabolism | Proteobacteria | NULL |
| 50% | gene_id_6359122 | 18 | 0.273 | oxc | K01577 | oxalyl-CoA decarboxylase |  | Glyoxylate and dicarboxylate metabolism | Proteobacteria | NULL |
| 15% | gene_id_665386 | 18 | 3.092 | Tas | K05275 | pyridoxine 4-dehydrogenase |  | Vitamin B6 metabolism | Actinobacteria | NULL |
| 50% | gene_id_665386 | 18 | 1.932 | Tas | K05275 | pyridoxine 4-dehydrogenase |  | Vitamin B6 metabolism | Actinobacteria | NULL |
| 15% | gene_id_8528059 | 18 | 0.018 | ich-Y | K18290 | itaconyl-CoA hydratase |  | C5-Branched dibasic acid metabolism | Proteobacteria | Bradyrhizobium |
| 50% | gene_id_8528059 | 18 | 0.038 | ich-Y | K18290 | itaconyl-CoA hydratase |  | C5-Branched dibasic acid metabolism | Proteobacteria | Bradyrhizobium |
| 15% | gene_id_9147865 | 18 | 0.004 | AKR1C3 | K04119 | aldo-keto reductase family 1 member C3 |  | Folate biosynthesis | NULL | NULL |
| 50% | gene_id_9147865 | 18 | 0.046 | AKR1C3 | K04119 | aldo-keto reductase family 1 member C3 |  | Folate biosynthesis | NULL | NULL |
| 15% | gene_id_9211047 | 18 | 0.951 | hypothetical | NULL | NULL | NULL | NULL | Actinobacteria | NULL |
| 50% | gene_id_9211047 | 18 | 0.644 | hypothetical | NULL | NULL | NULL | NULL | Actinobacteria | NULL |
| 15% | gene_id_9760539 | 18 | 0.043 | hypothetical | NULL | NULL | NULL | NULL | Proteobacteria | Bradyrhizobium |
| 50% | gene_id_9760539 | 18 | 0.149 | hypothetical | NULL | NULL | NULL | NULL | Proteobacteria | Bradyrhizobium |
| 15% | gene_id_9760540 | 18 | 0.118 | ptsI | K08483 | phosphoenolpyruvate-protein phosphotransferase (PTS system enzyme I) |  | Transporters | Proteobacteria | Bradyrhizobium |
| 50% | gene_id_9760540 | 18 | 0.350 | ptsI | K08483 | phosphoenolpyruvate-protein phosphotransferase (PTS system enzyme I) |  | Transporters | Proteobacteria | Bradyrhizobium |
| 15% | gene_id_9924387 | 18 | 0.199 | faaH | K16171 | fumarylacetoacetate (FAA) hydrolase | Tyrosine degradation, tyrosine => homogentisate | Tyrosine metabolism | Actinobacteria | NULL |
| 50% | gene_id_9924387 | 18 | 0.130 | faaH | K16171 | fumarylacetoacetate (FAA) hydrolase | Tyrosine degradation, tyrosine => homogentisate | Tyrosine metabolism | Actinobacteria | NULL |

**Table S3:**
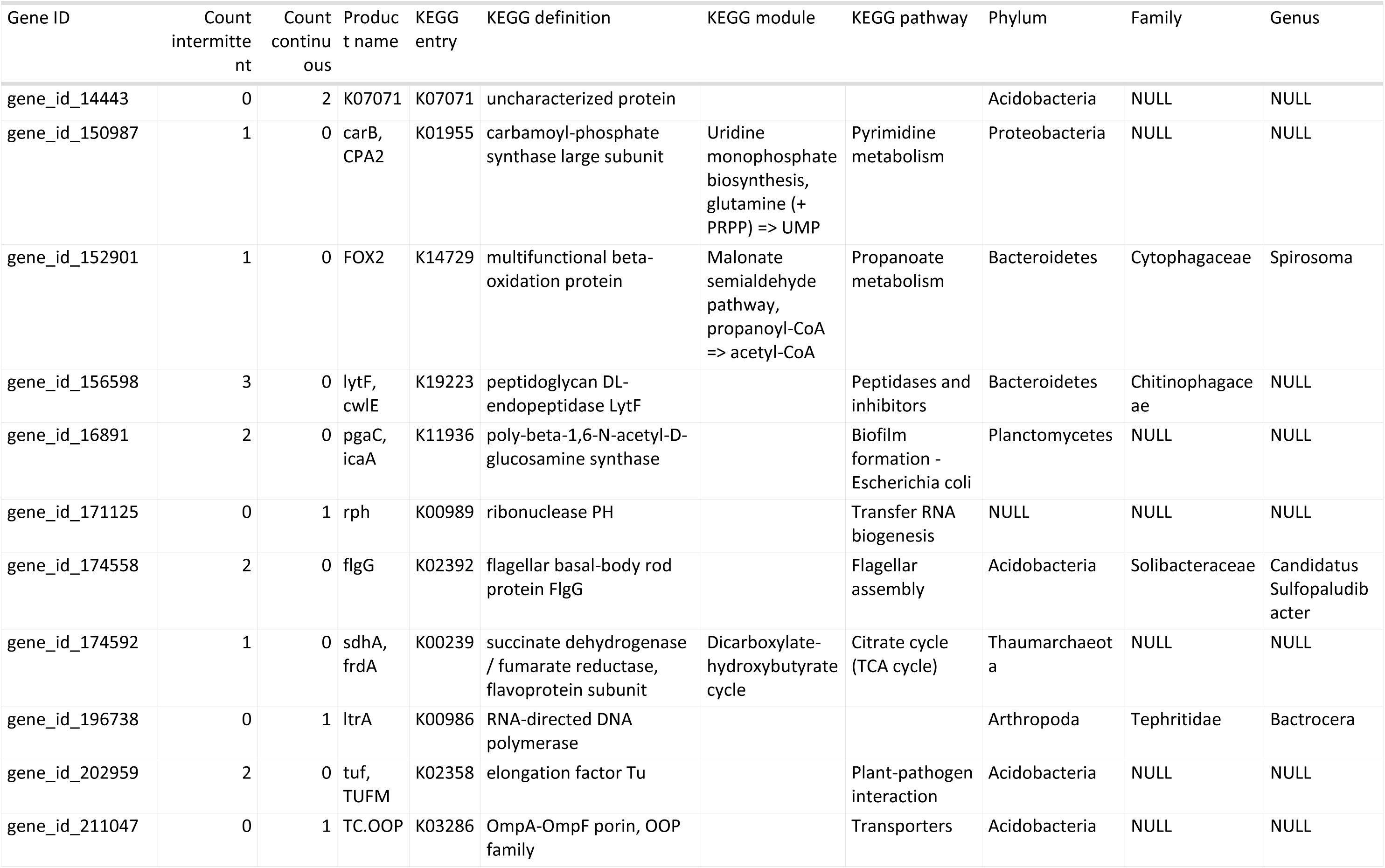
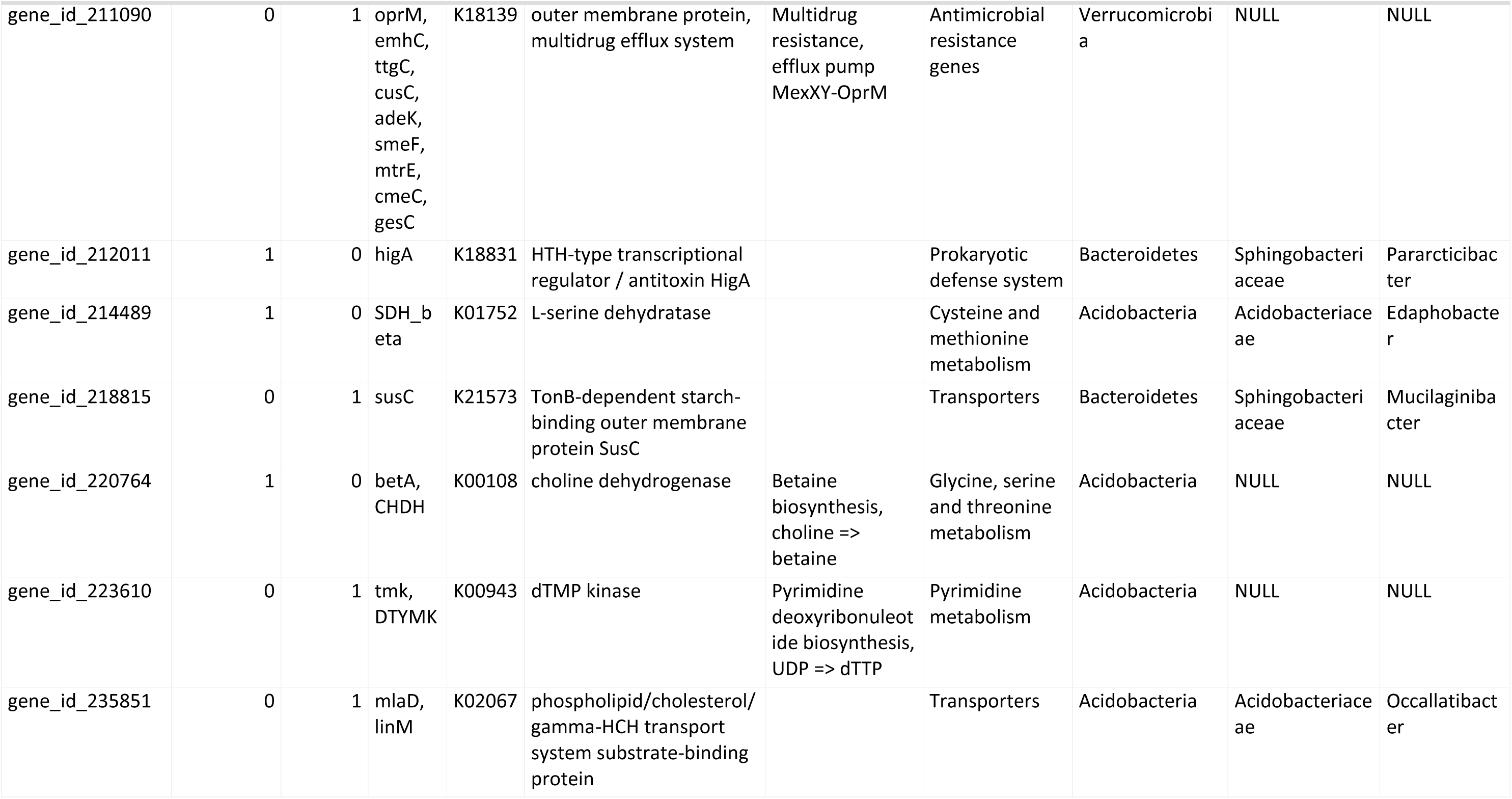
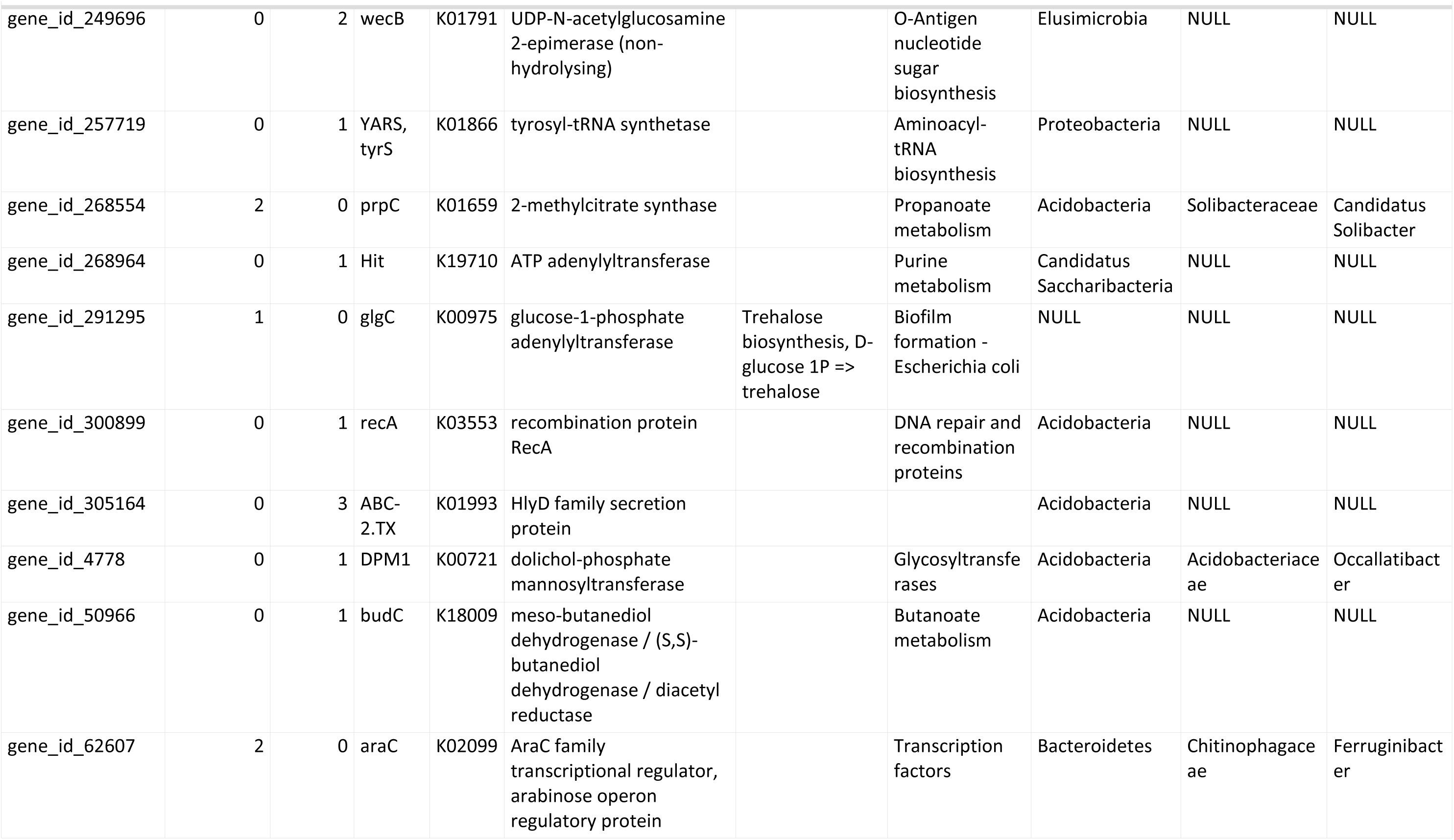
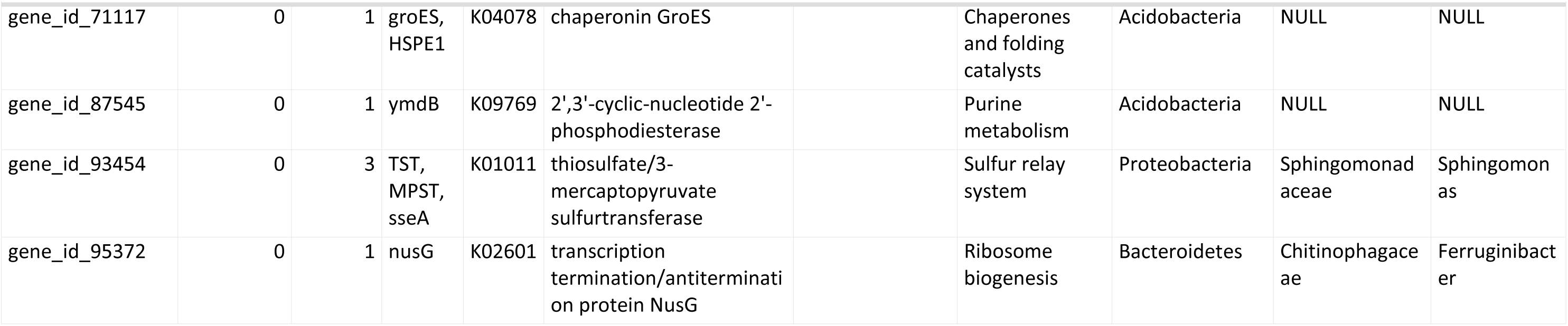
Potentially LGT genes overrepresented in inoculated samples.

| Gene ID | Count<br>intermitte<br>nt | Count<br>continu<br>ous | Produc<br>t name | KEGG<br>entry | KEGG definition | KEGG module | KEGG pathway | Phylum | Family | Genus |
| --- | --- | --- | --- | --- | --- | --- | --- | --- | --- | --- |
| gene_id_14443 | 0 | 2 | K07071 | K07071 | uncharacterized protein |  |  | Acidobacteria | NULL | NULL |
| gene_id_150987 | 1 | 0 | carB,<br>CPA2 | K01955 | carbamoyl-phosphate<br>synthase large subunit | Uridine<br>monophosphate<br>biosynthesis,<br>glutamine (+<br>PRPP) => UMP | Pyrimidine<br>metabolism | Proteobacteria | NULL | NULL |
| gene_id_152901 | 1 | 0 | FOX2 | K14729 | multifunctional beta-<br>oxidation protein | Malonate<br>semialdehyde<br>pathway,<br>propanoyl-CoA<br>=> acetyl-CoA | Propanoate<br>metabolism | Bacteroidetes | Cytophagaceae | Spirosoma |
| gene_id_156598 | 3 | 0 | lytF,<br>cwIE | K19223 | peptidoglycan DL-endopeptidase LytF |  | Peptidases and<br>inhibitors | Bacteroidetes | Chitinophagace<br>ae | NULL |
| gene_id_16891 | 2 | 0 | pgaC,<br>icaA | K11936 | poly-beta-1,6-N-acetyl-D-glucosamine synthase |  | Biofilm<br>formation -<br>Escherichia coli | Planctomycetes | NULL | NULL |
| gene_id_171125 | 0 | 1 | rph | K00989 | ribonuclease PH |  | Transfer RNA<br>biogenesis | NULL | NULL | NULL |
| gene_id_174558 | 2 | 0 | flgG | K02392 | flagellar basal-body rod<br>protein FlgG |  | Flagellar<br>assembly | Acidobacteria | Solibacteraceae | Candidatus<br>Sulfopaludib<br>acter |
| gene_id_174592 | 1 | 0 | sdhA,<br>frdA | K00239 | succinate dehydrogenase<br>/ fumarate reductase,<br>flavoprotein subunit | Dicarboxylate-<br>hydroxybutyrate<br>cycle | Citrate cycle<br>(TCA cycle) | Thaumarchaeot<br>a | NULL | NULL |
| gene_id_196738 | 0 | 1 | ltrA | K00986 | RNA-directed DNA<br>polymerase |  |  | Arthropoda | Tephritidae | Bactrocera |
| gene_id_202959 | 2 | 0 | tuf,<br>TUFM | K02358 | elongation factor Tu |  | Plant-pathogen<br>interaction | Acidobacteria | NULL | NULL |
| gene_id_211047 | 0 | 1 | TC.OOP | K03286 | OmpA-OmpF porin, OOP<br>family |  | Transporters | Acidobacteria | NULL | NULL |
| gene_id_211090 | 0 | 1 | oprM,<br>emhC,<br>ttgC,<br>cusC,<br>adeK,<br>smeF,<br>mtrE,<br>cmeC,<br>gesC | K18139 | outer membrane protein,<br>multidrug efflux system | Multidrug<br>resistance,<br>efflux pump<br>MexXY-OprM | Antimicrobial<br>resistance<br>genes | Verrucomicrobi<br>a | NULL | NULL |
| gene_id_212011 | 1 | 0 | higA | K18831 | HTH-type transcriptional<br>regulator / antitoxin HigA |  | Prokaryotic<br>defense system | Bacteroidetes | Sphingobacteri<br>aceae | Pararcticibac<br>ter |
| gene_id_214489 | 1 | 0 | SDH_b<br>eta | K01752 | L-serine dehydratase |  | Cysteine and<br>methionine<br>metabolism | Acidobacteria | Acidobacteriace<br>ae | Edaphobacte<br>r |
| gene_id_218815 | 0 | 1 | susC | K21573 | TonB-dependent starch-<br>binding outer membrane<br>protein SusC |  | Transporters | Bacteroidetes | Sphingobacteri<br>aceae | Mucilaginiba<br>cter |
| gene_id_220764 | 1 | 0 | betA,<br>CHDH | K00108 | choline dehydrogenase | Betaine<br>biosynthesis,<br>choline =><br>betaine | Glycine, serine<br>and threonine<br>metabolism | Acidobacteria | NULL | NULL |
| gene_id_223610 | 0 | 1 | tmk,<br>DTYMK | K00943 | dTMP kinase | Pyrimidine<br>deoxyribonucleot<br>ide biosynthesis,<br>UDP => dTTP | Pyrimidine<br>metabolism | Acidobacteria | NULL | NULL |
| gene_id_235851 | 0 | 1 | mIaD,<br>linM | K02067 | phospholipid/cholesterol/<br>gamma-HCH transport<br>system substrate-binding<br>protein |  | Transporters | Acidobacteria | Acidobacteriace<br>ae | Occallatibact<br>er |
| gene_id_249696 | 0 | 2 | wecB | K01791 | UDP-N-acetylglucosamine<br>2-epimerase (non-<br>hydrolysing) |  | O-Antigen<br>nucleotide<br>sugar<br>biosynthesis | Elusimicrobia | NULL | NULL |
| gene_id_257719 | 0 | 1 | YARS,<br>tyrS | K01866 | tyrosyl-tRNA synthetase |  | Aminoacyl-<br>tRNA<br>biosynthesis | Proteobacteria | NULL | NULL |
| gene_id_268554 | 2 | 0 | prpC | K01659 | 2-methylcitrate synthase |  | Propanoate<br>metabolism | Acidobacteria | Solibacteraceae | Candidatus<br>Solibacter |
| gene_id_268964 | 0 | 1 | Hit | K19710 | ATP adenyllyltransferase |  | Purine<br>metabolism | Candidatus<br>Saccharibacteria | NULL | NULL |
| gene_id_291295 | 1 | 0 | glgC | K00975 | glucose-1-phosphate<br>adenyllyltransferase | Trehalose<br>biosynthesis, D-glucose 1P =><br>trehalose | Biofilm<br>formation -<br>Escherichia coli | NULL | NULL | NULL |
| gene_id_300899 | 0 | 1 | recA | K03553 | recombination protein<br>RecA |  | DNA repair and<br>recombination<br>proteins | Acidobacteria | NULL | NULL |
| gene_id_305164 | 0 | 3 | ABC-2.TX | K01993 | HlyD family secretion<br>protein |  |  | Acidobacteria | NULL | NULL |
| gene_id_4778 | 0 | 1 | DPM1 | K00721 | dolichol-phosphate<br>mannosyltransferase |  | Glycosyltransfe<br>rases | Acidobacteria | Acidobacteriace<br>ae | Occallatibact<br>er |
| gene_id_50966 | 0 | 1 | budC | K18009 | meso-butanediol<br>dehydrogenase / (S,S)-<br>butanediol<br>dehydrogenase / diacetyl<br>reductase |  | Butanoate<br>metabolism | Acidobacteria | NULL | NULL |
| gene_id_62607 | 2 | 0 | araC | K02099 | AraC family<br>transcriptional regulator,<br>arabinose operon<br>regulatory protein |  | Transcription<br>factors | Bacteroidetes | Chitinophagace<br>ae | Ferruginibact<br>er |

| Gene ID | Count<br>intermitte<br>nt | Count<br>contin<br>ous | Produc<br>t name | KEGG<br>entry | KEGG definition | KEGG module | KEGG pathway | Phylum | Family | Genus |
| --- | --- | --- | --- | --- | --- | --- | --- | --- | --- | --- |
| gene_id_71117 | 0 | 1 | groES,<br>HSPE1 | K04078 | chaperonin GroES |  | Chaperones<br>and folding<br>catalysts | Acidobacteria | NULL | NULL |
| gene_id_87545 | 0 | 1 | ymdB | K09769 | 2',3'-cyclic-nucleotide 2'-<br>phosphodiesterase |  | Purine<br>metabolism | Acidobacteria | NULL | NULL |
| gene_id_93454 | 0 | 3 | TST,<br>MPST,<br>sseA | K01011 | thiosulfate/3-mercaptopyruvate<br>sulfurtransferase |  | Sulfur relay<br>system | Proteobacteria | Sphingomonad<br>aceae | Sphingomon<br>as |
| gene_id_95372 | 0 | 1 | nusG | K02601 | transcription<br>termination/antiterminati<br>on protein NusG |  | Ribosome<br>biogenesis | Bacteroidetes | Chitinophagace<br>ae | Ferruginibact<br>er |

## Notes

### Competing Interest Statement

The authors have declared no competing interest.

